# Branching, crosslinking and decentralization of microtubules accelerates intracellular assembly

**DOI:** 10.1101/2025.06.05.658035

**Authors:** Apurba Sarkar, Alex Mogilner, Raja Paul

## Abstract

Before cell division, mitotic spindle is assembled from chromosomes and centrosomes. After the cell division, Golgi organelles assemble from multiple vesicles scattered across daughter cells. These are but two examples of intracellular assembly of vesicles, organelles and chromosomes made possible by dynamic microtubules. The most prominent microtubule networks are centrosome-focused asters that ‘search’ for the vesicles and chromosomes, but there are also microtubules originating from the vesicles and chromosomes, raising the question whether a coordination between multiple microtubule networks optimizes the assembly process. This study uses a computational model to examine how microtubule dynamics influence the assembly of organelles from vesicles. The model includes two microtubule populations: microtubules anchored to the vesicles, which drive local clustering, and ‘central’ microtubules anchored to the centrosome that aggregate the vesicles globally. Simulations show that a microtubule decentralization – balanced contribution from both microtubule populations — accelerates the assembly of tens of vesicles, but that assigning all microtubules to hundreds of vesicles optimizes the assembly. Directionally biased microtubule growth, particularly when avoiding spontaneous catastrophe events, further accelerates the assembly. Additionally, microtubule branching, when occurring at optimal angles and spacings, enhances the assembly’s efficiency. Lastly, rapid crosslinking of overlapping central and ‘local’ microtubules can drastically accelerate the assembly. Applying this model to the spindle assembly in early mitosis reveals similar insights. The model suggests that the observed multiple microtubule networks optimize the intracellular assembly processes when molecular resources are limited.

**SIGNIFICANCE:** Assembly of intracellular structures is a time-sensitive process driven by multiple dynamic microtubule networks. For example, vesicles must be aggregated by microtubules to form Golgi apparatus after cell division, while mitotic spindle assembly requires bringing chromosomes closer together by microtubules growing from spindle poles and chromosomes. Delays in these processes can lead to genomic instability and disease. Using modeling, we show how microtubules originating from centrosomes, vesicles, or chromosomes can be optimally distributed to minimize the assembly time. The model reveals roles of microtubule decentralization, branching and crosslinking in accelerating assembly. The model suggests that the intracellular assembly can be optimized by diversifying microtubule networks and enhancing their interactions.

## INTRODUCTION

Assembly of organelles and other subcellular structures, i.e. fusion of mitochondria (1), is a crucial part of cell dynamics. Several such processes – reassembly of Golgi in daughter cells at the end of cell division (2), aggregation of melanosomes (pigment granules) in fish melanophore cells (3), and congression of chromosomes to mitotic spindle in prometaphase (4) – are driven by microtubules (MTs) and dynein motors. For example, to divide Golgi complex between daughter cells during cell division, Golgi stacks fragment into tens to thousands of vesicles that disperse stochastically across the cytoplasm in mitosis (5–7). As the mother cell transitions into telophase, these vesicles are gathered at centrosomes of the daughter cell, where the vesicles reassemble into Golgi stacks and ribbons (2).

This reassembly crucially depends on dynamic instability of MTs (8, 9), in which MT plus ends grow outward, then undergo a ‘catastrophe’, shrink inward, toward the MT organizing center (in this case centrosome), where MT minus ends are anchored, and then resume the growth after a ‘rescue’. The catastrophes and rescues occur with certain frequencies at random times, and the resulting stochastic process effectively enables the MT plus ends to explore the cell volume. Namely, in an elementary act of the ‘search-and-capture’ process, a MT plus end growing in a random direction encounters a Golgi vesicle by sheer chance, dynein motors coating the vesicles transport it to the centrosome (10). A similar mechanism governs the central aggregation of pigment granules in melanophore cells (11). Last, but not least, accurate chromosome segregation during mitosis (12, 13) depends on mitotic spindle assembly that starts in early prometaphase with connecting chromosomes to two MT asters centered at two centrosomes. (Some cells opt for an alternative, centrosome-independent, spindle assembly pathway (14).) The early search-and-capture models posited that these connections are made by random contacts between the growing MT plus ends and kinetochores, small proteinacious structures on the centromeres of the chromosomes (15).

The central, global search-and-capture process is usually complemented by a decentralized, local assembly made possible by nucleation and anchoring of MTs not only at the centrosomes, but also at the Golgi membranes (16) and pigment granules (11). MTs are also nucleated near and associate with kinetochores in complex and understudied ways (13, 17– 20). Specifically, a non-centrosomal MT growing from one Golgi vesicle randomly encounters another such vesicle after which dynein motors on the second vesicle transport it toward the MT minus end on the first one, ending in merger of the two vesicles (10, 21). Similarly, melanophore pigment granules nucleate MTs (11, 22) that search randomly the cellular space, capture other pigment granules and bring the granules together in the dynein-dependent way (3, 11). While central and local aggregation processes could be sufficient on their own, they usually cooperate accelerating the assembly (3, 10, 11). This raises the broad question: *Is there an optimal strategy of the combined central-local MT-dynein-driven assembly, and if so, do cells in fact optimize the assembly?*

Furthermore, while chromosomes do not appear to aggregate locally, there is growing evidence that mitotic spindle assembly relies on interactions of long centrosomal MTs with short kinetochore MTs (23, 24). Namely, the direct kinetochore capture by centrosomal MTs is not the principal pathway of the spindle assembly, but rather either centrosomal MTs capture minus ends of kinetochore-associated short MTs, or the kinetochore-MTs capture the centrosomal MT (4), after which bundles of the kinetochore-MTs are pulled toward the centrosomes in a dynein-dependent way (17–20, 25, 26). Absence of the kinetochore-MTs impedes the spindle assembly (27). Thus, like organelle assembly, spindle formation may also rely on a combined central-local search to optimize the spindle assembly.

The global and local searches in the spindle are further optimized by several factors, most prominently by making the searches less random and biasing MT growth towards chromosomes. The best understood relevant pathway is mediated by spatial gradients of RanGTP protein around chromosomes; this gradient increases capture probability by stabilizing MTs growth near chromosomes, preventing catastrophe before capture (28–30). Additional factor that can accelerates the search is random chromosome movement during early prometaphase (31).

Centrosomal MTs and MTs anchored to organelles, vesicles and chromosomes are not the only MT networks in the cell. MTs are also nucleated at and branch from the sides of pre-existing MTs in a process mediated by Augmin and other proteins (23, 24, 32–34). This branching MT nucleation is considered to be the major source of spindle MTs (35), and acentrosomal spindles rely mainly on the branched MT architecture to self-assemble (36). In centrosomal spindles, MT branching is a part of the spindle formation: inhibiting centrosomal MTs increases MT branching near chromosomes, while disrupting Augmin-dependent MT formation enhances centrosomal MT nucleation (37, 38). This suggests not only cooperation among different MT networks in the assembly processes, but also that this cooperation proceeds under constrants of limited molecular resources, i.e. of a finite supply of tubulin (37, 38). This raises the second key question: *What is the optimal coordination between several MT networks for the subcellular assemblies?*

In this study, we use a computational model to simulate the assembly of cellular organelles and vesicles (vesicles hereafter) through a MT-driven search process achieved by both central searcher MTs, originating from a single centrosome, and local searcher MTs produced by the vesicles. We quantify the roles of these two populations and examine potential contributions of MT branching, spatial and angular biases of MT growth, and MT crosslinking. We then apply a similar model to simulate chromosomal capture process in early mitosis. Our model conserves the total number of MTs in the cell, reflecting the limited supply of tubulin (37, 38). We find that purely local, decentralized assembly gives the fastest result for hundreds of vesicles, while the combined central/local search is optimal for tens of vesicles (the latter is also relevant for the spindle assembly). The simulations also reveal that unregulated MT branching does not improve the assembly, however, spatially and angularly biased accelerates the process. The model suggests that random movements of vesicles or chromosomes and/or rapid inter-MT crosslinking can drastically accelerate the assembly. The simulations result in quantitative estimates that agree with reported experimental measurements.

## METHODS

### Vesicle Assembly Model

We examine the assembly of vesicles to a central location in the cell through a *search and capture* mechanism mediated by dynamic MTs within a three-dimensional spherical cell, as illustrated in Figure 1 *a*. The system includes a centrally located centrosome and *N*_ves_ vesicles, initially randomly dispersed and modeled as small spheres of radius *R*_ves_, diffusing within the cell. Both vesicles and centrosome are capable of nucleating MTs. Initially, all vesicles contribute equally to the pool of *N*_vMT_ vesicle-derived MTs, such that each vesicle grows *N*_vMT_ / *N*_ves_ MTs. These MTs, along with *N*_cMT_ centrosome-derived MTs, are engaged in dynamic instability and explore the cellular space (see Supporting Material for simulation details). When a vesicle is reached by an MT originating from another vesicle, the two instantly merge to a position near either of the original vesicles; the merged vesicle’s volume is equal to the sum of the volumes of the pair before the merger. The merged vesicle nucleates a number of MTs equal to the sum of MT numbers nucleated by the merged pair. Newly formed MTs grow in random directions from the merged vesicle. If a vesicle is captured by a centrosomal MT, it is transported toward the cell center instantly and contributes its MTs to the central searcher. MTs that reach the cell boundary undergo immediate depolymerization, resetting their search.

**FIGURE 1:**
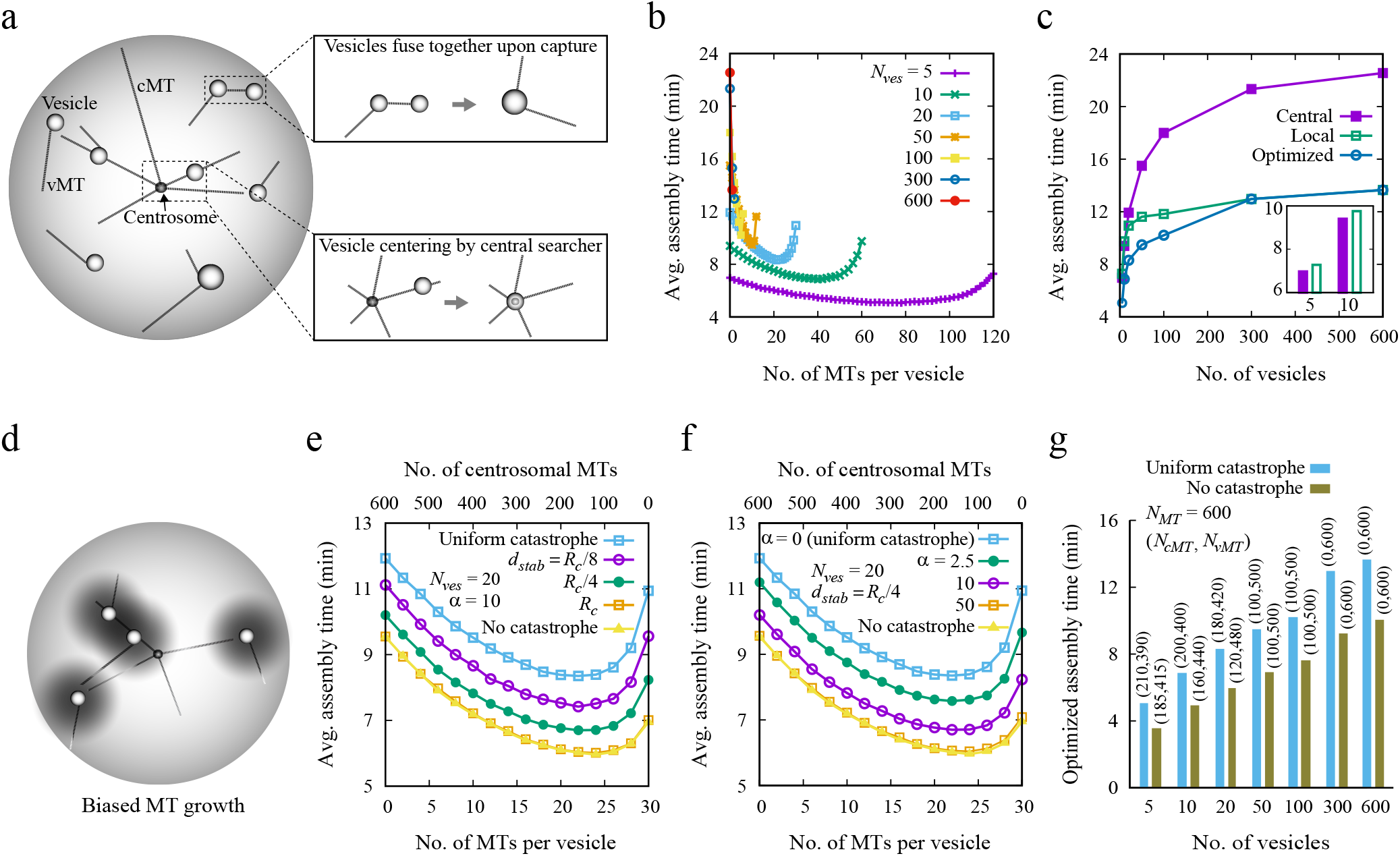
Efficient vesicle assembly with distinct MT populations. (*a*) Schematic of the assembly process. Central and local searcher MTs originate from the centrosome and vesicles, respectively. Vesicles merge upon contact via local MTs, conserving volume and MT number (top *inset*). Central searcher MTs guide vesicles toward the center (bottom *inset*). (*b*) Mean assembly time vs. number of MTs per vesicle for *N*_ves_ = 5, 10, 20, 50, 100, 300, and 600 with uniform catastrophe frequency *f*_*c*_ = 0.016 sec^−1^. (*c*) Assembly time vs. vesicle number with MTs assigned only to central or local searchers. The optimal curve combines both MT types, derived from the minima in (*b*). *Inset*: for few vesicles, using only central searcher MTs leads to slightly reduced search times. (*d*) Vesicle assembly with MT bias under stabilizing gradients. Stabilizing agents decay exponentially around vesicles (shaded regions), enhancing MT growth toward targets and effectively destabilizing growth away. (*e*–*f*) Assembly time vs. MTs per vesicle for *N*_ves_ = 20 under biased conditions: (*e*) varying *d*_stab_ at fixed *α* = 10; (*f*) varying *α* at *d*_stab_ = *R*_*c*_/ 4. Comparisons include unbiased MTs with *f*_*c*_ = 0.016 sec^−1^ (uniform catastrophe) and *f*_*c*_ = 0 (no spontaneous catastrophe). The top *x*-axis shows corresponding centrosomal MT counts. (*g*) Optimized assembly time vs. vesicle number for both catastrophe scenarios. For 600 total MTs, optimal allocations to centrosomal (*N*_cMT_) and vesicle MTs (*N*_vMT_) are labeled above bars.

We consider the following scenarios during the vesicle search process:

### Unbiased search without branching

MTs explore the cellular space without any bias and branching and with unaltered dynamic instability parameters (growth velocity, *v*_*g*_, shrinkage velocity, *v*_*s*_, catastrophe frequency, *f*_*c*_, and rescue frequency, *f*_*r*_).

### Biased search modulated by a local stabilizing gradient

A spatial bias is introduced by incorporating a stabilizing gradient centered around each vesicle, which influences MT dynamics by modulating the catastrophe frequency. Inspired by the role of RanGTP in chromosomal spindle assembly, this bias is implemented by modeling catastrophe frequency as an exponentially decreasing function of stabilizing agent concentration, which decays with distance from the vesicle (see Eq. S1, Eq. S2, and Supporting Material). This effectively biases MT growth toward nearby vesicles (see Figure 1 *d*). We also explore cases where spontaneous catastrophes are suppressed entirely due to a uniformly distributed global stabilizing agent.

### MT branching

MTs can form branches from existing filaments. Each MT is allowed to generate a fixed number (*N*_*br*_) of branch MTs at a rate *k*_*br*_, originating at different distances along the mother MT (see Figure 2 *a*). Secondary branching is excluded. If a branch MT shrinks back to its nucleation site, it dissociates and can re-emerge later from the growing mother MT. Similarly, if the mother MT shrinks past the branch origin, the branch detaches immediately. To conserve the total number of MTs, the nucleation of a branch MT from a central (or local) MT leads to the removal of a randomly chosen MT from the same population. Conversely, when a branch dissociates, a new MT immediately grows from the original source of its mother MT.

**FIGURE 2:**
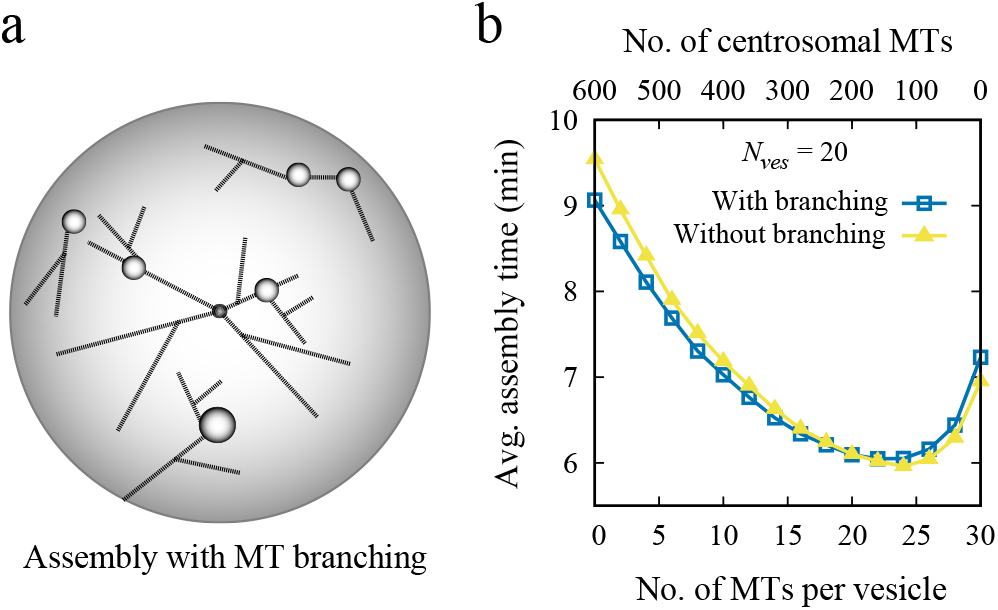
Minimal effect of unrestricted MT branching on vesicle assembly time. (*a*) Schematic of MT branching: each MT can produce branches at a given rate, preserving the total number of MTs. Branches form at arbitrary angles and distances along the mother MT. (*b*) Average assembly time vs. number of MTs per vesicle for *N*_ves_ = 20, with and without branching. MTs grow without spontaneous catastrophe (*f*_*c*_ = 0). The top *x*-axis indicates corresponding centrosomal MT counts.

Further details of the simulation model are provided in Supporting Material, and parameter values are listed in Table S1.

### Chromosomal Assembly Model

We simulated the *search and capture* of chromosomes by MTs using an agent-based model (15, 31, 39). The system includes dynamic components – chromosomes, kinetochores, and MTs - and two static centrosomes positioned at opposite poles of a spherical cell. Each centrosome serves as an anchoring site for the minus ends of *N*_*cMT*_ centrosomal MTs. Chromosomes are modeled as cylindrical rods with random orientations and are uniformly distributed throughout the cellular volume. They move randomly (on the scale of tens of seconds, they diffuse) while maintaining self-avoidance, preventing physical overlap. In addition to translational motion, chromosomes are subject to small rotational displacements at each computational step to represent thermal fluctuations in orientation. Each chromosome contains two sister kinetochores, represented as cylindrical elements positioned back-to-back on the chromosome surface, located at the midpoint along its length. From each kinetochore, *N*_*kt MT*_ kinetochore-associated MTs (ktMTs) emerge and undergo dynamic instability during the search process. Centrosomal MTs can capture chromosomes either through direct contact with kinetochores or via interaction with ktMTs, with the latter occurring at a rate *k*_*inter*-*MT*_. Once a kinetochore establishes a connection to a centrosome, either directly through a centrosomal MT or indirectly via ktMTs, it is considered captured and is no longer available for further capture by centrosomal MTs. Notably, the translational and rotational motions of a chromosome continue even after one or both kinetochores have been captured. The *search and capture* process is complete once all kinetochores are successfully attached to spindle MTs.

MTs begin to shrink immediately after contacting the cell boundary or colliding with a chromosome arm. Upon successful attachment to their target, the captured MTs are replaced by new ones originating from the centrosomes or kinetochores, ensuring that the total number of searcher MTs remains constant throughout the simulation. A schematic representation of the chromosomal assembly process is shown in Figure 5 *a*. Further technical details, including model implementation and parameter values, are provided in Supporting Material and Table S1. As described earlier in the vesicle assembly model, here we also explored three scenarios: (a) MT search with finite catastrophe rates, without bias or branching; (b) inclusion of a RanGTP-like spatial bias, and a comparison with a condition where no spontaneous catastrophes occur inside the cell; and (c) MT branching from both centrosomal and kinetochore MTs.

To explore the parameter space, we performed 2,000 - 20,000 simulations for each parameter set using different initial configurations. The model was implemented in C and run on an Intel(R) Xeon(R) CPU with a clock speed of 2.20 GHz and 50 GB of RAM. Depending on complexity of the simulated scenario, each simulation took from a few seconds to several minutes. Data analysis and visualization were conducted using GNUPLOT and MATLAB (The MathWorks, Natick, MA).

## RESULTS

### Coordination among the central and local searching MTs optimizes the vesicle assembly

We first address the central question: how does the assembly time depend on the MT numbers allocated to the central (centrosomal) and/or local (vesicular) asters if the total MT number is a constant? To find the answer, we examined how the average assembly time varies with the number of MTs emanating from each initial vesicle (Figure 1 *b*). In the simulations, the catastrophe frequency was set so that MTs could grow, on average, up to 1.5 times the cell radius before starting to shorten.

We found that when the number of vesicles is small (roughly less than 20), allocating all MTs to the central aster, or all MTs to the vesicles, leads to almost the same assembly times, with slightly faster assembly in the central search compared to purely local search (Figure 1, *b* and *c*). The intuitive way to understand this result is based on a simple estimate of the average search time for *n* targets by an *N*-MT aster. This time scales as ~ ln (*n*)/*N* (28), because the search time decreases inversely proportionally to the number of searching MTs (two searchers find a target twice faster than one searcher) – this theoretical result agrees with our simulations (compare Figure 1 *c* with Figure S1, *a* and *b*). The logarithmic dependence on the number of targets, apparent from the simulation results (Figure 1 *c*), is due to the fact that the search for *n* targets is over when all *n* targets are captured independently. The probability of capturing one target increases with time as (1 exp(−*t*/ *τ*)), where *τ* is characteristic time of a single capture; the probability of capturing *n* targets independently is (1 exp(−*t* /*τ*))^*n*^. After characteristic time of capturing all *n* targets, *T*, the probability (1 − exp (−*T*/ *τ*)) ^*n*^ is less than 1 by a small number ϵ; as *T* > *τ*, (1 − exp (−*T*/ *τ*)) ^*n*^ ≈ 1 − *n* · exp (−*T* /*τ*) = 1 − ϵ, thus, *T* = *τ* ln (*n* / ϵ). This argument suggests that one central searcher with *N* MTs captures *n* vesicles after time ~*t*_0_ ln (*n*)/ *N*, where *t*_0_ is a constant parameter. The same argument says that if all MTs are distributed equally between *n* vesicles, than one of the vesicles captures (*n* − 1) others after time ~*t*_0_ ln (*n* − 1) (*N*/ *n*). As *any* of the vesicles can capture any other one, all vesicles would be assembled after time ~ (1 /*n*) · *t*_0_ ln (*n* − 1) (*N* / *n*) ≈ *t*_0_ ln (*n*) /*N* – similar to the time of the central assembly. The slight advantage of the central search in this case stems from the geometry: the most distant vesicles from the cell center are one cell radius away, while two most distant vesicles are two radii apart.

The central result of the simulations is that in the case of up to ~ 100 initial vesicles, the average assembly time reaches a minimum at a nontrivial optimal partitioning of MTs between a minority (~20 − 30%) MT number assigned to the centrosome and the rest equally divided between the initial vesicles (Figure 1, *b* and *g*). The time gain is modest but nontrivial and can be understood as follows: initially, some vesicles are much closer to each other than to the cell center, and allowing the vesicles to merge locally accelerates the assembly. At the later stage of the search, when many vesicles are already captured by the centrosome, the number of central MTs becomes the majority, and the last stage of the search is effectively the central one, which is relatively fast for the small number of remaining vesicles.

Another nontrivial result is that for a great number of vesicles, ≳ 300, the fastest assembly corresponds to purely local search, and time will only be lost if any MT are assigned to the central search. The explanation is that when the initial vesicle number is great, capturing proximal neighbors is a very fast process, then the merged vesicles with accumulating MT number capture the next neighbors, and so on, with the ‘cloud’ of vesicles effectively coarsening. This also effectively shifts the vesicles from the cell periphery inward (the peripheral vesicles on average merge inward, toward multiple neighbors in the cell interior. Interior vesicles capture peripheral ones more effectively by searching uniformly in all directions, unlike peripheral vesicles that often search unproductively toward the cell edge), leading to their assembly at an interior position away from the cell center (Figure S1 *c*). Thus, bringing the merged vesicle to the center likely requires at least a few central MTs. Lastly, note that for the great initial vesicle number, the local search is roughly twice faster than the central search (Figure 1 *c*), which is due to smaller, on average, distances between the vesicles capturing each other than between the cell center and vesicles.

We also found that at low vesicle numbers, a very small catastrophe frequency can significantly speed up assembly, even when all MTs are vesicle-nucleated (Figure S1 *d*). At higher catastrophe rates, local MTs from distant vesicles may not persist long enough to reach their targets, while lower catastrophe frequencies enhance MTs persistently grow toward the targets, improving capture efficiency. The simulations also demonstrated that when the vesicles move randomly in the cell, greater effective diffusion coefficient leads to faster assembly (Figure S1 *e*). When the diffusion is faster than certain threshold, differences between various central, local and optimal assembly scenario disappears, because rapid movements bring a vesicle close to the searcher faster than dynamically unstable MTs can capture this vesicle. In what follows, we simulate initial vesicles with a small diffusion constant of the order of ~ 0.001 μm^2^ /sec (other than during merger events), which keeps MT partitioning strategies relevant for efficient assembly.

### Spatially biased MT dynamics further accelerates the assembly

Several studies suggest that spatially biased MT growth toward targets enhances search efficiency (28–31). In early mitosis, chromosomes establish a localized biochemical environment that promotes MT nucleation and directional growth. Among the mechanisms responsible, the steep gradient of RanGTP, a

RAS-related nuclear protein, centered around chromosomes plays a key role in biasing MT dynamics (23). Within this gradient, MTs tend to grow steadily toward chromosomes while undergoing rapid collapse when growing away. This directional bias can be mimicked by modulating the catastrophe frequency in response to local stabilizing signals (28).

While RanGTP’s role is well studied in mitotic chromosome assembly, we investigate whether a similar gradient centered around individual vesicles could bias MTs and reduce the time required for the vesicle assembly. The bias is implemented by reducing the catastrophe frequency near vesicles, modeled as an exponential decay controlled by a stabilizing gradient centered on each vesicle (Figure 1 *d*). Two parameters define the influence of this gradient: the spatial range *d*_*stab*_ (how far the stabilizing effect extends from a vesicle), and sensitivity *α* (how much the catastrophe frequency is decreased by the concentration of the stabilizing agent) (see Eq. S1, Eq. S2, and Supporting Material).

For both central and local searches, biased MT dynamics consistently resulted in faster assembly compared to the unbiased MT dynamics (Figure S1 *f*). The optimized hybrid central/local assembly also showed an improved performance with bias. Moreover, increasing either of the parameters, *d*_*stab*_ or *α*, further shortened the assembly time (Figure 1, *e* and *f*). At large values of both *d*_*stab*_ and *α*, the system approaches a limit where the catastrophe frequency within the cell becomes negligible – similar to an idealized case where MTs never undergo a catastrophe inside the entire cell (Figure 1, *e* and *f*). In this regime, no spontaneous MT catastrophe within the cell occurs, other than when an MT hits the cell boundary. Under such conditions, assigning all MTs to vesicles minimizes the assembly time regardless of the number of the vesicles (Figure S1 *g*). However, when the catastrophe rate is finite, allocating all MTs to the central searcher may become more efficient at low vesicle counts (Figure S1, *d* and *g)*.

Overall, optimal combinations of central and local searchers lead to the fastest assembly in both biased and unbiased scenarios. With very few vesicles, most MTs should be centrally located, while with many vesicles, assigning nearly all MTs to local searchers is optimal (Figure 1 *g*). The lowest assembly times occur when spontaneous catastrophes are absent. Notably, this trend remains consistent even when the total number of MTs is halved or doubled (Figure S1, *a* and *b*). In the following, we simulate MTs without spontaneous catastrophes inside the cell, experiencing catastrophes only at the cell boundary.

### Effects of MT branching on the assembly process

As multiple reports indicate that MT branching could play a crucial role in subcellular processes (reviewed in (40)), we explored whether branching from existing MTs could affect vesicle assembly. In the model, the total number of MTs remains constant before and after the branch nucleation (Figure 2 *a*; see also METHODS). In Figure 2 *b*, we compare assembly times with and without MT branching. Each growing central or local searcher MT was allowed to generate up to two branches at a rate of 0.1 sec^−1^, with branches forming at random angles and distances from the mother MT’s nucleation site (Figure 2). Note that the literature reports a wide range of branching angles, including values close to 0^°^, around 20^°^ to 60^°^, near 90^°^, and close to 180^°^ (32, 34, 41–44) (also reviewed in (33, 40)). We found that such random branching did not significantly alter the assembly time compared to the no-branching case (Figure 2 *b*).

We next explored whether branching could reduce the assembly time by enhancing MT coverage in peripheral regions of the cell where central MTs splay apart. Two promising strategies would be: (a) initiating branches only after the mother MT grows beyond a certain distance from its origin (defined by the *branching length, l*_*br*_), and (b) limiting the orientation of a branch to a specific angle range relative to the mother MT (the *branch angle, θ*_*br*_). To evaluate the impact of these two factors, we plotted the computed average assembly time as a function of parameters *l*_*br*_ and *θ*_*br*_ for three cases: (i) all MTs originating from the centrosome (*N*_*cMT*_ = 600), (ii) all MTs assigned to the vesicles (*N*_*vMT*_ = 600), and (iii) an optimized central/local search uncovered in previous non-branching simulations (Figure 3, *a*–*c*; see also Figure 2 *b*).

**FIGURE 3:**
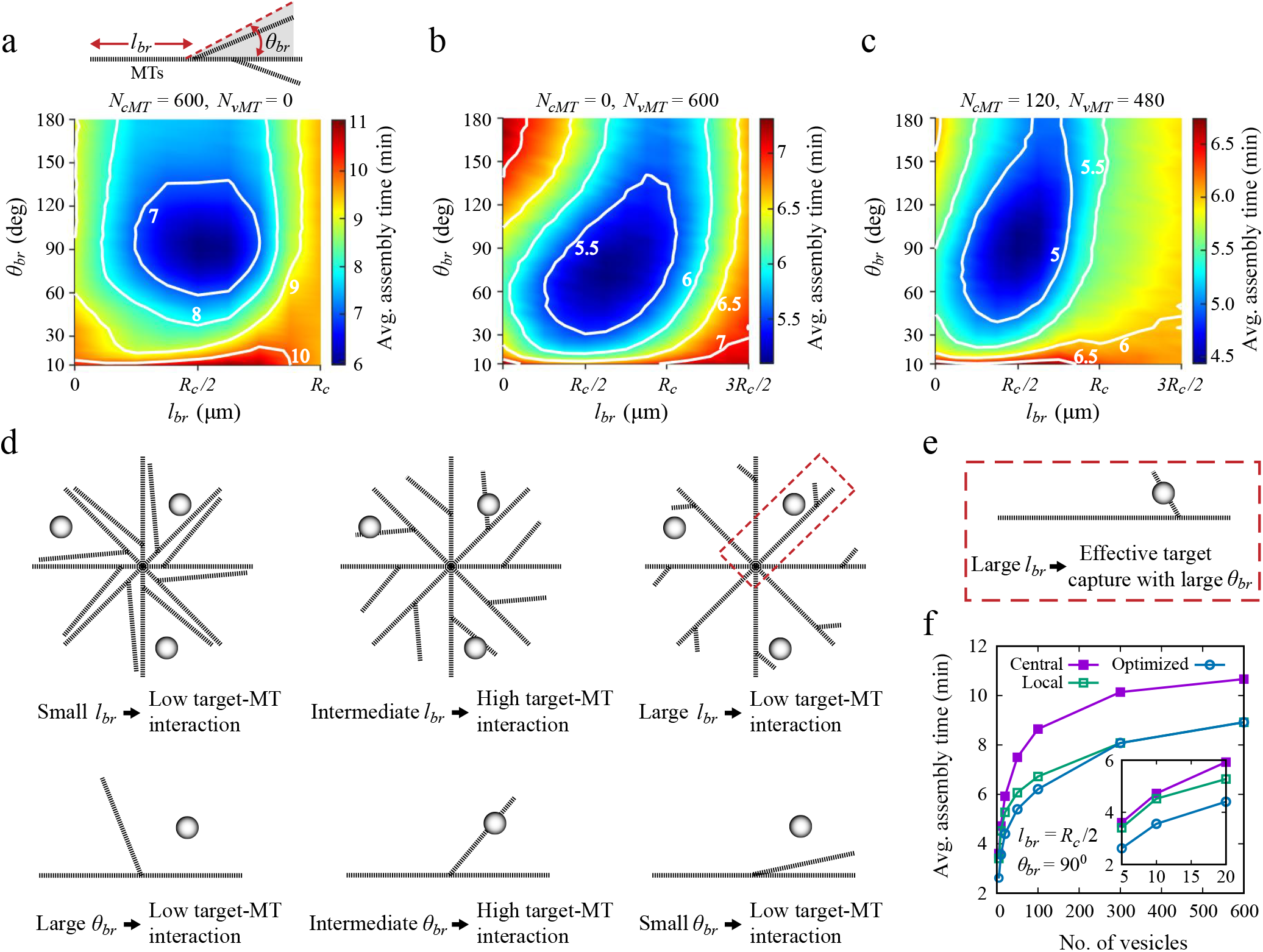
Optimizing vesicle assembly with MT branching. (*a*) Average assembly time as a function of branch length (*l*_br_) and angle (*θ*_br_) for *N*_ves_ = 20, with all MTs as central searchers (*N*_cMT_ = 600). (*b*) Same as (*a*), but with all MTs as local searchers (*N*_vMT_ = 600). (*c*) Assembly time for an optimized mix of central and local searchers (*N*_cMT_ = 120, *N*_vMT_ = 480), minimizing time without branching (see Figure 2 *b*). (*d*) Schematic showing how intermediate *l*_br_ and *θ*_br_ enhance MT-vesicle contacts. (*e*) At larger *l*_br_, wider branching angles improve MT-vesicle targeting compared to narrower ones (see red dotted box in (*d*)). (*f*) Average assembly time vs. vesicle number for *l*_br_ = *R*_*c*_ / 2 and *θ*_br_ = 90^°^, comparing central, local, and optimized MT assignments.

Because vesicle-nucleated MTs can grow up to about 2*R*_*c*_ in length (for a vesicle located near the cell boundary), while centrosomal MTs’ length is limited by the cell radius, we explored a broader range of *l*_*br*_ for vesicle MT-based configurations (Figure 3, *b* and *c*) and limited the branching length range to 0–*R*_*c*_ in the central MT-only scenario (Figure 3 *a*). Intermediate values of both *l*_*br*_ and *θ*_*br*_ produced shorter assembly times than in cases without branching. When central MTs were used, the minimum search time occurred around *l*_*br*_ ≈ *R*_*c*_/ 2 (±1.5) μm and *θ*_*br*_ 90^°^ ≈ (±30^°^) (Figure 3, *a* and *c*). When all MTs were assigned to the vesicles, the optimal region shifted slightly to *l*_*br*_ 2*R*_*c*_ 3 2 μm and *θ*_*br*_ ≈ 75^°^ (±30^°^) (Figure 3 *b*). The reason for this shift likely stems from the broader spatial distribution of vesicles, allowing vesicular MTs to probe farther regions of the cell compared to central MTs.

These optimal values emerge from a balance between too-early and too-late branching (Figure 3 *d*). If *l*_*br*_ is too small, branching occurs near the MT aster center, spreading MTs too soon and diluting density where it is most needed. Conversely, if branching occurs too late, the mother MTs may have already reached sparse regions, limiting the chances for the branches to capture nearby vesicles. Similarly, extreme values of *θ*_*br*_ are suboptimal: large angles effectively return daughter MTs to regions already searched by mother MTs, while MTs branched at small angles search too close to mother MTs (Figure 3 *d*). Intermediate angles allows branches to cover the cell space more effectively, improving the capture of vesicles that might otherwise be missed. Notably, at higher *l*_*br*_, wider *θ*_*br*_ values can, in fact, improve efficiency by enabling branches to target gaps between diverging mother MTs (Figure 3, *d* and *e*).

When parameter *l*_*br*_ approaches ~ 0.9*R*_*c*_ in the central-only case, the effect of branching becomes negligible, irrespective of *θ*_*br*_ values (Figure 3 *a*). Here, mother MTs undergo catastrophe soon after branching, causing branches to disappear before contributing meaningfully and resulting in assembly times similar to the non-branching scenario. In contrast, the vesicle-only system benefits from slightly longer *l*_*br*_ and intermediate *θ*_*br*_, likely due to its ability to reach farther into the cell. The hybrid configuration, which includes both central and vesicular MTs for *N*_*ves*_ = 20, yields an optimal region similar to the central-only case (Figure 3 *c*; see also Figure 3 *a*), but overall assembly time is minimized when all MTs are allocated to the vesicles at higher vesicle counts, especially with *l*_*br*_ ≈ *R*_*c*_/2 and *θ*_*br*_ ≈ 90^°^ (Figure 3 *f*).

### Sensitivity of assembly time to the number of branch MTs, rate of branch nucleation, and initial vesicle distribution

To evaluate how sensitive the assembly process is to model parameters, we varied a few parameters at a time while keeping the other parameters fixed at baseline values (Table S1). We first explored how the assembly time depends on the branching length (*l*_*br*_) and branch angle (*θ*_*br*_) for different numbers of MT branches (*N*_*br*_ = 2, 4, 8) (Figure S2, *a*–*f*). In both search scenarios (MTs assigned either to the central searcher or to the vesicles), increasing *N*_*br*_ expanded the range of *l*_*br*_ and *θ*_*br*_, yielding shorter assembly times. More branches increased the chances of reaching the target, either by steering some branches in its direction or by covering more space. For a broader range of *l*_*br*_, additional branches improved spatial coverage (Figure S2 *g*). Even at large values of *θ*_*br*_, some MTs nucleated at favorable angles, maintaining efficiency (Figure S2 *h*). However, at very small values of *θ*_*br*_, the search space narrowed, and adding more branches in that confined region reduced efficiency, increasing assembly time.

We then examined the effect of the branching rate (*k*_*br*_ = 0.1, 0.01, 0.001 sec^−1^), assuming that each MT could form two branches (Figure S3). At a high rate of 0.1 sec^−1^, branches typically nucleated shortly after the mother MT reached *l*_*br*_, yielding optimal *l*_*br*_ values around half the cell radius (Figure S3, *a* and *d*). At lower rates, delays in nucleation meant that branching often occurred beyond *l*_*br*_, making those values less effective. Smaller *l*_*br*_ values helped initiate earlier branching and improved outcomes at intermediate rates (Figure S3, *b* and *e*). However, at the lowest rate, branching was too infrequent to support efficient search, and assembly times remained high across the full *l*_*br*_ range (Figure S3, *c* and *f*).

We also studied how initial vesicle distribution influences the assembly time by restricting the vesicles to a narrow annular region 0.2*R*_*c*_ (2 μm) near the cell edge (Figure S4 *a*). Assigning all MTs to vesicles produced assembly time trends similar to that in the case of uniformly distributed vesicles (Figure S4 *b*; see also Figure 3 *b*). In contrast, when all MTs originated from the central searcher, the assembly time increased near *l*_*br*_ ≈ 0.8*R*_*c*_, just before the vesicle-rich region (Figure S4 *c*). This resulted from the model’s rule that forming a new branch removes an existing central MT, so early branching near *l*_*br*_ ≈ 0.8*R*_*c*_ reduced the number of long MTs reaching the vesicles (Figure S4 *d*). Initiating branches earlier (*l*_*br*_ ≈ *R*_*c*_ / 2) or closer to the cell edge (*l*_*br*_ ≈ *R*_*c*_) improved targeting, as mother MTs either directed branches toward vesicles or reached them directly. However, assembly time is lower with branch nucleation near *l*_*br*_ ≈ *R*_*c*_ /2 than closure to the cell edge.

We further investigated how the vesicle number influences assembly with the peripheral vesicle distribution. When all MTs were distributed among the vesicles – a configuration favoring optimized assembly time – increasing the number of vesicles slightly increased assembly time, likely due to the broader spatial distribution of the targets. However, when all MTs originated from the central searcher, more vesicles, paradoxically, led to a significant decrease in assembly time (Figure S4 *e*). In this case, once a vesicle near the periphery was captured, it was transported toward the center and became a new site for MT nucleation. This process amplified the central searcher’s reach by generating additional MTs, which in turn improved the likelihood of capturing remaining vesicles. Thus, with more vesicles near the periphery, the feedback between vesicle capture and enhanced MT nucleation at the center accelerated and improved assembly.

### Strong inter-MT coupling diminishes the impact of MT branching

So far, we have examined the assembly in which the capture occurs through direct MT–vesicle contact. However, the following scenario is possible: two MTs originating at different vesicles intersect in the cytoplasm, which could lead to MT-MT crosslinking by MT-based molecular motors and/or protein crosslinkers, which indirectly connects the two vesicles together. Then, the motors can, in principle, rapidly bring the vesicles together leading to the merger. There is a relevant example of a similar process in mitotic spindle formation, where centrosomal and kinetochore-derived MTs get interconnected by dynein and NuMA resulting in formation of MT bundles that link kinetochores to the spindle pole (20, 26). Inspired by this example, we examined what happens if we include such crosslinking interactions between MTs from different sources, either central or vesicular, and their branches. We assume in the model that such interactions lead to either vesicle merging or transport to the center, depending on MT origin (Figure 4 *a*). For simplicity, we assumed that upon MT-MT intersection (specifically, when shortest MT-MT distance is less than a threshold equal to MT diameter), the merger of the origins of the MT pair occurs at a single rate *k*_inter-MT_.

**FIGURE 4:**
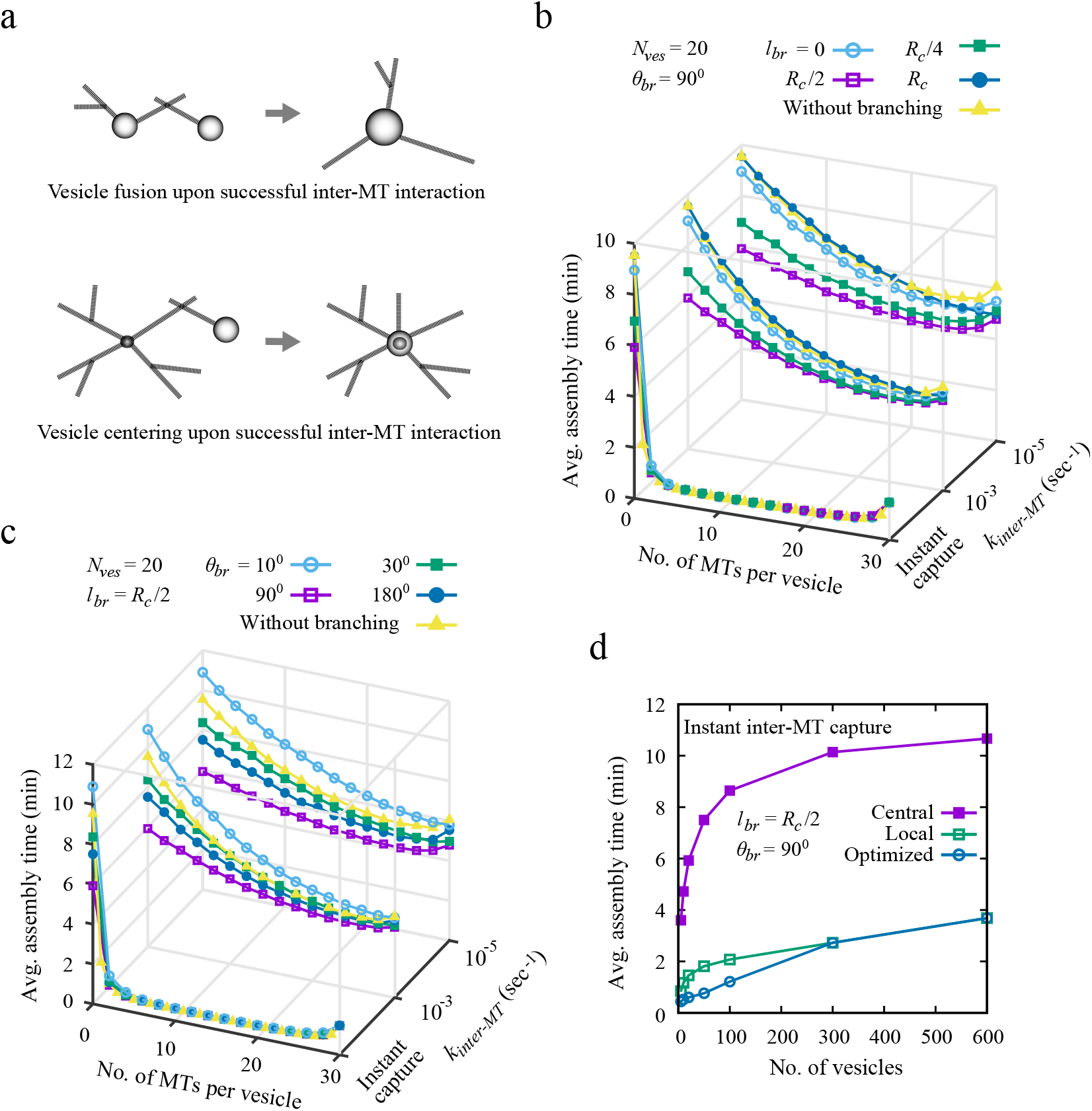
Strong inter-MT coupling diminishes branching effects and accelerates assembly, regardless of limited local searchers. (*a*) Schematic of vesicle assembly showing fusion or central translocation following inter-MT interaction. (*b*) Average assembly time vs. MTs per vesicle for *N*_ves_ = 20 at *θ*_br_ = 90^°^ and varying branch lengths (*l*_br_ = 0, *R*_*c*_ / 4, *R*_*c*_ / 2, *R*_*c*_), for different inter-MT interaction rates: *k*_inter-MT_ = 10^−5^, 10^−3^ sec^−1^, and instantaneous capture. (*c*) Assembly time for fixed *l*_br_ = *R*_*c*_ / 2 and branch angles *θ*_br_ = 10^°^, 30^°^, 90^°^, 180^°^. Results in (*b*–*c*) are compared with the no-branching case. (*d*) Assembly time vs. vesicle number for *l*_br_ = *R*_*c*_ / 2 and *θ*_br_ = 90^°^ under instantaneous inter-MT capture, comparing central, local, and optimized MT distributions.

**FIGURE 5:**
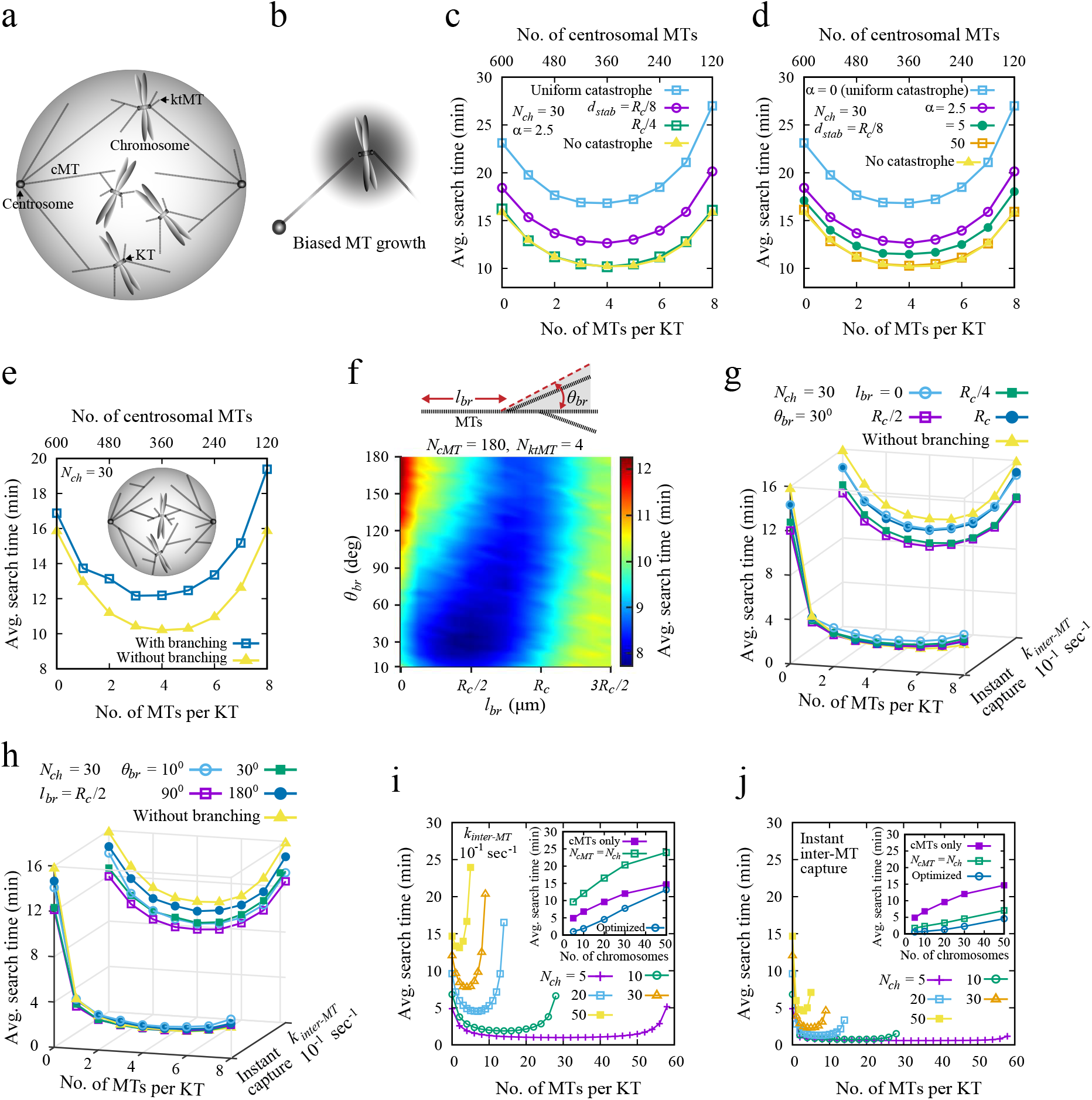
Efficient chromosomal assembly across MT configurations. (*a*) Schematic of chromosome capture during mitosis, via either centrosomal MTs or associated kinetochore MTs (ktMTs). (*b*) Biased MT growth under a stabilizing RanGTP gradient, which decays exponentially around chromosomes (dark shaded regions), promotes MTs growing toward chromosomes and effectively destabilizes those growing away. (*c*–*d*) Assembly time vs. MTs per KT for biased models with *N*_ch_ = 30 (*N*_kt_ = 60), varying (*c*) *d*_stab_ (*α* = 2.5) and (*d*) *α* (*d*_stab_ = *R*_*c*_ / 8). Results are compared with uniform catastrophe (*f*_*c*_ = 0.016 sec^−1^) and no catastrophe inside cell (*f*_*c*_ = 0). Top *x*-axis shows corresponding centrosomal MT numbers. (*e*) Search time vs. MTs per KT with and without branching (*f*_*c*_ = 0). *Insets* show MT branching during chromosome search, with branches emerging at variable distances and angles. (*f*) Search time vs. *l*_br_ and *θ*_br_ using an optimized MT setup (*N*_cMT_ = 180, *N*_ktMT_ = 4), based on minimum search time without branching (Figure 5 *e*). (*g*–*h*) Search time vs. MTs per KT at *k*_inter-MT_ = 10^−1^ sec^−1^ and with instantaneous MT capture: (*g*) *θ*_br_ = 30^°^ and *l*_br_ = 0, *R*_*c*_ / 4, *R*_*c*_ / 2, *R*_*c*_; (*h*) *l*_br_ = *R*_*c*_ / 2 and *θ*_br_ = 10^°^, 30^°^, 90^°^, 180^°^. Both compared to no-branching cases. (*i*–*j*) Search time vs. MTs per KT for *N*_ch_ = 5, 10, 20, 30, 50 under *k*_inter-MT_ = 10^−1^ sec^−1^ (*i*) and instantaneous capture (*j*). *Insets* show search time vs. chromosome number with either centrosome-only MTs (300 cMTs per centrosome) or minimal MT allocation across centrosomes (*N*_*cMT*_ = *N*_*ch*_). Optimal search times are taken from the minima in (*i*) and (*j*).

To evaluate the role of this potential process, we varied the number of local searcher MTs per vesicle and measured average assembly times for different interaction rates *k*_inter-MT_, with and without branching (Figure 4, *b* and *c*). We first fixed the branching angle at *θ*_*br*_ = 90^°^ and varied the branching distance *l*_*br*_, allowing up to two branches per MT to nucleate at a rate of 0.1 sec^−1^. At low rate *k*_inter-MT_, the results were similar to those in the non-MT-interacting cases: moderate *l*_*br*_ values improved the assembly, while small or large *l*_*br*_ slowed it down. As rate *k*_inter-MT_ increased, differences between the assembly times for different *l*_*br*_ values diminished, and at high interaction rates, all conditions converged to similar assembly times, especially when more local MTs were present. Under instantaneous inter-MT capture, the assembly took place super-fast, because hundreds of MTs effectively packed the intracellular space allowing almost instant interactions between the vesicles. In that case, branching offered no added benefit, as vesicle interactions occurred too quickly for branching to influence search efficiency. We also fixed parameter *l*_*br*_ = *R*_*c*_ / 2 and varied parameter *θ*_*br*_. Consistent with earlier results, the assembly was fastest around *θ*_*br*_ ~ 90^°^ at low values of *k*_inter-MT_, while both smaller and greater angles slowed the process. Again, higher interaction rates reduced these differences, and under instant inter-MT capture, assembly times across all angles became nearly identical to those without branching, making the specific angle of MT branching less critical. Moreover, when MTs frequently interact, interactions among vesicles via local searcher MTs strongly dominate the assembly process over central MT-only configurations, significantly reducing assembly times across vesicle counts (Figure 4 *d*; see also Figure 3 *f*). During spindle assembly, the MT-MT crosslinking seems to take mere tens of seconds (4, 20); thus, based on our simulations, the MT-MT crosslinking with correspondingly high rates can significantly accelerate the assembly.

### Chromosome capture during spindle formation is optimized under conditions similar to those of the vesicle assembly

Similar to the vesicle assembly, spindle formation during mitosis relies on two MT populations: centrosomal MTs from two opposite poles and kinetochore MTs from each chromosome (4, 20, 27). Chromosomes can be captured, in principle, either directly via interactions between associated kinetochores and centrosomal MTs, or indirectly through inter-MT interactions between centrosomal and kinetochore MTs at a rate of ~ 0.1 sec^−1^ (4) (Figure 5 *a*).

Our simulations showed, first, that a spatial gradient of MT-stabilizing factors, like RanGTP, around chromosomes, speeds up the search by reducing catastrophe events in MTs growing toward chromosomes (Figure 5, *b*–*d*). This effect depends on parameter *α*, which controls the MT dynamic sensitivity to RanGTP, and parameter *d*_*stab*_, the stabilization range (see Eq. S1 and Eq. S2; Supporting Material). Under strong MT stabilization conditions (higher *α* and/or *d*_*stab*_), average search times approach those in systems without spontaneous MT catastrophes inside the cell (Figure 5, *c* and *d*).

We next examined the potential role of the MT branching in the chromosome capture, if the total MT number is constant. In these simulations, catastrophes occur only upon encountering obstacles such as the cell wall or chromosome arms. MT branches emerge at rate 0.1 sec^−1^ once the mother MT reaches a length *l*_*br*_ and within an angle *θ*_*br*_ relative to the mother (Figure 5, *insets* in *e* and *f*). Similar to the case of the vesicle assembly, unrestricted branching (*θ*_*br*_ = 180^°^, *l*_*br*_ = 0) offers no benefit over the non-branching scenario (Figure 5 *e*). To identify favorable conditions, we varied parameters *l*_*br*_ and *θ*_*br*_ using the MT arrangement that minimized the search time without branching (Figure 5 *f*; see Figure 5 *e* for the optimized MT combination). Optimal assembly is observed when branching occurs at *l*_*br*_ ≈ *R*_*c*_ / (±2 2.5 μm) and at branch angles *θ*_*br*_ ≈ 30^°^ (±15^°^) (Figure 5 *f*). This dynamics likely compensates for the reduced MT density near the spindle poles (conserving total MT number means reducing mother MT number when daughter MTs emerge) by initiating branches farther away from the poles, where mother MTs splay apart. Compared to the vesicle assembly model, the optimal parameter value *θ*_*br*_ is lower in the spindle case, likely due to spatial constraints: centrosomal MTs in the spindle assembly are directed inward from the opposite poles, so smaller branching angles help focus growth toward chromosomes. However, excessively small angles delay the spindle assembly.

We next estimate the assembly time as a function of the number of kinetochore MTs, allowing instant inter-MT interactions to enable rapid KT capture upon intersection of centrosomal and kinetochore MTs. We compared this scenario to that with a finite MT interaction rate (0.1 sec^−1^), previously used in Figure 5, *c*–*f*. With *θ*_*br*_ = 30^°^ and varying *l*_*br*_, finite MT interaction rate shortens search time by branching, but instant capture upon MT-MT interaction yields similar times with or without branching (Figure 5 *g*). Likewise, changing *θ*_*br*_ at fixed *l*_*br*_ = *R*_*c*_ / 2 under the instant capture condition has a minimal effect (Figure 5 *h*), suggesting that the MT branching is less important when inter-MT interactions are fast enough.

Finally, we explore how the assembly time varies with the number of MTs per kinetochore for different chromosome numbers, considering both finite inter-MT interaction rate (*k*_*inter* − *MT*_ = 0.1 sec^−1^) and instantaneous capture upon intersection of centrosomal and kinetochore MT pair (Figure 5, *i* and *j*). We increase the number of kinetochore MTs while keeping the total MT number fixed, ensuring that each centrosome has at least *N*_*ch*_ centrosomal MTs for a system with *N*_*ch*_ chromosomes. Since each chromosome has two kinetochores, at least *N*_*ch*_ centrosomal MTs per pole are needed to connect all chromosomes to the spindle. We observe that the search time reaches a minimum at certain optimal number of kinetochore MTs (Figure 5 *i*). However, when centrosomal MTs are limited to *N*_*cMT*_ = *N*_*ch*_, assembly is slower compared to a configuration where all MTs are centrosomal (Figure 5 *i, inset*). In this case, the lower availability of centrosomal MTs, combined with finite inter-MT interactions, delays chromosome capture. By contrast, under instantaneous inter-MT interaction, kinetochore MTs significantly improve efficiency for any chromosome number, even when centrosomal MTs are scarce (Figure 5 *j*; see also *inset*).

## DISCUSSION AND CONCLUSIONS

The main conclusion of our study, as summarized in Figure 6, is that when MT pool is limited, there is an optimal combination of central (centrosome-anchored) and local (vesicle-anchored) MTs for the fastest assembly of tens of Golgi vesicles or pigment particles at the cell center. The optimal MT partitioning corresponds to a minor fraction of MTs (on the order of 20 to 30%) assigned initially to the centrosome, while the majority of the MTs should be equipartitioned between the vesicles. As the assembly proceeds, the centralized MT number will grow. The qualitative explanation for these predictions is that the local assembly far away from the cell center is much more effective than the centralized search by splayed-out MTs near the cell periphery, while the central search of few aggregates at the last stage accelerates the final assembly. The model predicts that in the characteristic mitotic spindles with tens of chromosomes, the fastest integration of all these chromosomes into the bipolar spindle occurs when roughly half (a few hundreds) of MTs emanate from two spindle poles, while the remainder – comprising several MTs per kinetochore – are associated with the chromosomes. These estimates semi-quantitatively (experimental measurements of MT numbers in live cells remain elusive) agree with observations of Golgi reassembly (10), pigment granules’ aggregations in melanophores (3, 11) and spindle assembly (20, 27).

**FIGURE 6:**
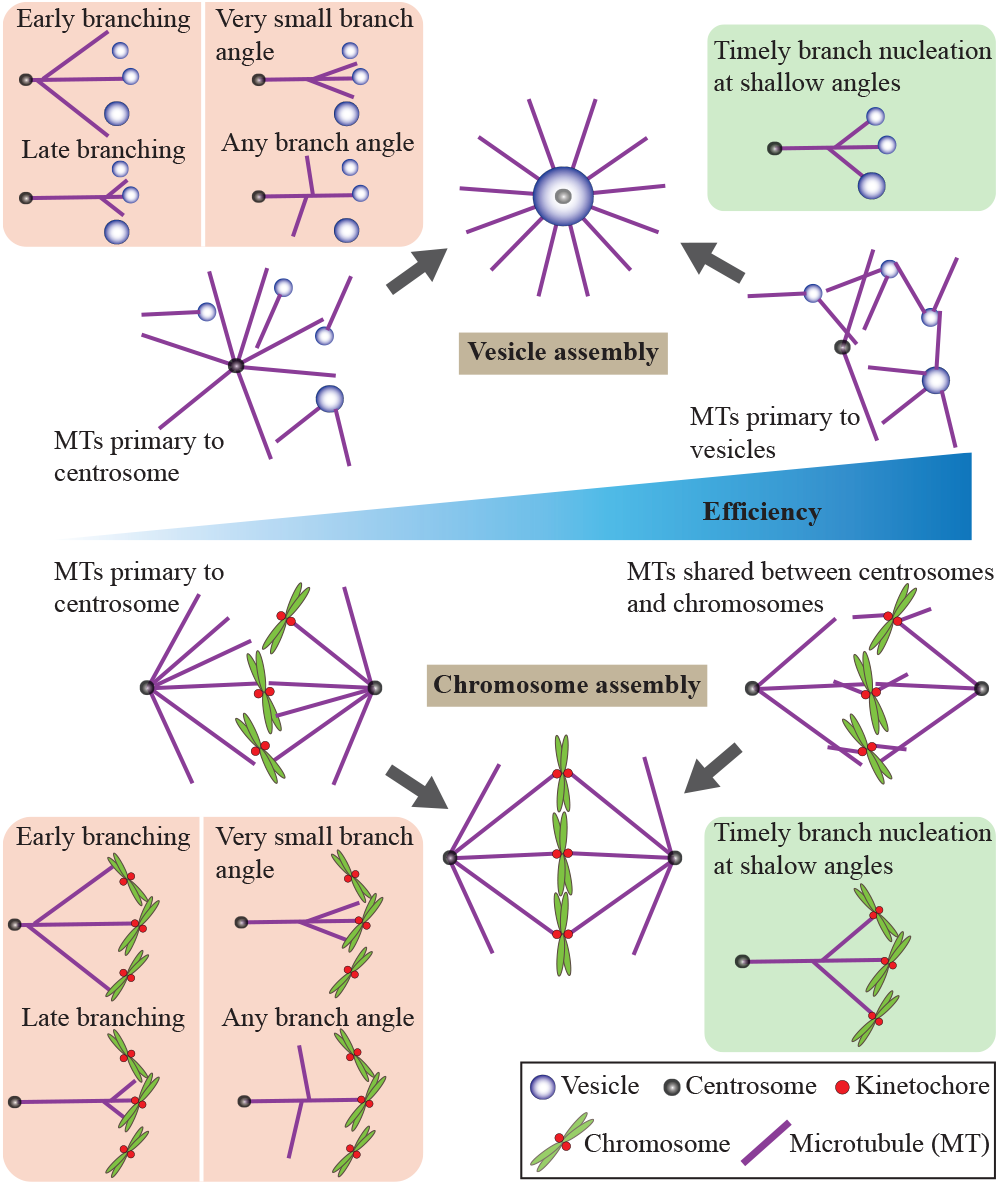
Summary of key outcomes showing efficient vesicle and chromosome assembly under different MT arrangements. Compared to cases where most MTs originate from centrosomes, assigning MTs primarily to vesicles or sharing them between centrosomes and chromosomes results in more efficient assembly. MT branches that nucleate either too early or too late, or at very narrow or wide angles, reduce search efficiency. In contrast, timely branching at intermediate distances and shallow angles improves overall assembly efficiency. Strong coupling between MTs from different origins reduces the effect of branching (not shown here; see Figure 4, *b* and *c*, and Figure 5, *g* and *h*).

Notably, the predicted acceleration in the optimal regime is relatively modest compared to purely local or centralized searches – just a few minutes faster – but at least in the spindle assembly case, every one of the gained minutes could be important (45). The model correctly estimates that Golgi vesicles can be reassembled in 10-20 minutes (10), while the spindle can be assembled in a few minutes (4, 27).

Interestingly, the model prediction for hundreds of vesicles or granules is that purely local search is the fastest - the local assembly of many proximal vesicles proves highly efficient and enables rapid assembly even in the absence of a centrosome. Indeed, Golgi-derived MTs can assemble the Golgi complex from dispersed ministacks without centrosomes (46), while melanosome aggregation in fish melanophores, can proceed via local MTs alone (11). The caveat of this prediction, however, is that the simulated vesicles may cluster away from the cell center, suggesting that a very few central MTs are likely required to gather the last few vesicle clumps to the centrosome.

The model suggests that the assembly is most effective in the absence of MT catastrophes inside the cell, away from the cell boundary. For finite catastrophe rates, shifting more MTs to local searches improves the outcome in general, while the optimal MT partitioning between the central and local searchers outlined in the previous paragraphs still holds. For finite catastrophe rates, the model predicts that a gradient of stabilizing agents around vesicles can accelerate the assembly. Similar prediction is made in the spindle assembly model, which aligns with theoretical (28, 31) and experimental findings (29, 30).

Our results show that unbiased MT branching does not improve the assembly, however, biased branching that occurs after the mother MT has grown to approximately half the cell radius, and when branches form at specific angles, around 90^°^ (±30^°^) for vesicle assembly and 30^°^ (±15^°^) for chromosome capture, significantly accelerates the assembly. The difference in the two optimal angles is due to the searching the entire cell volume during the vesicle assembly process, compared to a searching a smaller volume around the cell equator from the poles in the spindle assembly process. These findings are consistent with experimental observations of shallow-angled MT branching in mitotic spindles and interphase plant cells (32–34), suggesting that cells regulate branch initiation position and angle to maintain high MT density at the periphery (47).

If vesicles or chromosomes are engaged in rapid enough random movements during the assembly processes, the assembly duration decreases and MT partitioning strategy is irrelevant, because these movements bring a vesicle or a chromosome close enough to a searcher fast enough. The simulations show that an effective diffusion coefficient of the order of 0.001 μm^2^ / sec is the threshold between the two regimes. Thermal diffusion of a 0.1 μm-radius vesicles in cytoplasm with viscosity 1000 times higher than that of water, according to Stockes’ formula, is on the order of 0.001 μm^2^ /sec (48), so our simulations of the assembly of low-mobility vesicles, which are larger than 0.1 μm in radius and of ~ 1μm-large chromosomes are relevant.

Spindle assembly proceeds in the presence of crosslinking of MT pairs originating from the spindle poles (centrosomes) and the kinetochores, mediated by proteins such as dynein and NuMA (20, 26). In the spindle, such inter-MT crosslinking can occur within tens of seconds (20). The model demonstrates that MT crosslinking that is so fast can drastically accelerate the spindle assembly; similar result is valid for the vesicle assembly. The explanation is that in the presence of the fast crosslinking MTs essentially search for each other, rather than solely targeting vesicles, and it could be much faster to interconnect two vesicles by intersecting MTs than to find the vesicle by a MT. Interestingly, the model predicts that rapid crosslinking makes purely central search worse than the local assembly for any number of vesicles.

Several factors, in addition to more complex cell geometry, may require refining our model. It remains to be tested whether cells maintain a constant total MT number. Assembly of Golgi complex involves more complexity than simply bringing the vesicles together: only the trans side of the Golgi supports MT tethering (49), and MT nucleation may be restricted to specific vesicle regions (49, 50). Centrosomal MT aster may have a special role in correct Golgi organization (10) making the central search indispensable even for the case of hundreds of vesicles. Similarly, chromosome orientation (31), kinetochore architecture (39) and initial geometry of chromosomes’ distribution across the cell (51), and chromosome size distribution (52) affect the spindle assembly making it much more complex and nuanced than in our model here. MT pivoting, which we did not consider, could further accelerate the assembly (53, 54). There are additional MT networks in some cells, i.e. those anchored on the nuclear envelope in myotubes (55) and the apical membrane in polarized epithelial cells (56), which are not accounted for in the model. Last, but not least, the model does not yet include error correction mechanisms that are likely to prolong the correct assembly (57–59).

## AUTHOR CONTRIBUTIONS

R.P., A.M., and A.S. conceived the study and wrote the paper together. R.P. and A.S. developed the numerical code and ran the simulations.

## ACKNOWLEDGMENTS

R.P. and A.S. thank Indian Association for the Cultivation of Science (IACS) for funding and computational facilities. A.M. acknowledges the support from National Science Foundation (Grant no. DMS1953430).

## SUPPORTING CITATIONS

References (60–62) appear in the Supporting Material.

## REFERENCES

1. Westermann, B., 2010. Mitochondrial fusion and fission in cell life and death. Nat. Rev. Mol. Cell Biol. 11:872–884.

2. Gaietta, G. M., B. N. G. Giepmans, T. J. Deerinck, W. B. Smith, L. Ngan, J. Llopis, S. R. Adams, R. Y. Tsien, and M. H. Ellisman, 2006. Golgi twins in late mitosis revealed by genetically encoded tags for live cell imaging and correlated electron microscopy. Proc. Natl. Acad. Sci. USA. 103:17777–17782.

3. Lomakin, A. J., P. Kraikivski, I. Semenova, K. Ikeda, I. Zaliapin, J. S. Tirnauer, A. Akhmanova, and V. Rodionov, 2011. Stimulation of the CLIP-170–dependent capture of membrane organelles by microtubules through fine tuning of microtubule assembly dynamics. Mol. Biol. Cell. 22:4029–4037.

4. Renda, F., C. Miles, I. Tikhonenko, R. Fisher, L. Carlini, T. M. Kapoor, A. Mogilner, and A. Khodjakov, 2022. Non-centrosomal microtubules at kinetochores promote rapid chromosome biorientation during mitosis in human cells. Curr. Biol. 32:1049–1063.

5. Warren, G., and W. Wickner, 1996. Organelle Inheritance. Cell. 84:395–400.

6. Colanzi, A., C. Suetterlin, and V. Malhotra, 2003. Cell-cycle-specific Golgi fragmentation: how and why? Curr. Opin. Cell Biol. 15:462–467.

7. Colanzi, A., C. Sutterlin, and V. Malhotra, 2003. RAF1-activated MEK1 is found on the Golgi apparatus in late prophase and is required for Golgi complex fragmentation in mitosis. J. Cell Biol. 161:27–32.

8. Alberts, B., A. Johnson, J. Lewis, M. Raff, K. Roberts, and P. Walter, 2002. Molecular biology of the cell. New York: Garland Science.

9. Kirschner, M., and T. Mitchison, 1986. Beyond self-assembly: From microtubules to morphogenesis. Cell. 45:329–342.

10. Vinogradova, T., R. Paul, A. D. Grimaldi, J. Lončarek, P. M. Miller, D. Yampolsky, V. Magidson, A. Khodjakov, A. Mogilner, and I. Kaverina, 2012. Concerted effort of centrosomal and Golgi-derived microtubules is required for proper Golgi complex assembly but not for maintenance. Mol. Biol. Cell. 23:820–833.

11. Malikov, V., E. N. Cytrynbaum, A. Kashina, A. Mogilner, and V. Rodionov, 2005. Centering of a radial microtubule array by translocation along microtubules spontaneously nucleated in the cytoplasm. Nat. Cell Biol. 7:1213–1218.

12. Tolić-Nørrelykke, I. M., 2008. Push-me-pull-you: how microtubules organize the cell interior. Eur. Biophys. J. 37:1271–1278.

13. Pavin, N., and I. M. Tolić-Nørrelykke, 2014. Swinging a sword: how microtubules search for their targets. Syst. Synth. Biol. 8:1103.

14. de Saint Phalle, B., and W. Sullivan, 1998. Spindle Assembly and Mitosis without Centrosomes in Parthenogenetic Sciara Embryos. J. Cell Biol. 141:1383–1391.

15. Heald, R., and A. Khodjakov, 2015. Thirty years of search and capture: The complex simplicity of mitotic spindle assembly. J. Cell Biol. 211:1103.

16. Chabin-Brion, K., J. Marceiller, F. Perez, C. Settegrana, A. Drechou, G. Durand, and C. Poüs, 2001. The Golgi complex is a microtubule-organizing organelle. Mol. Biol. Cell. 12:2047–2060.

17. Maiato, H., C. L. Rieder, and A. Khodjakov, 2004. Kinetochore-driven formation of kinetochore fibers contributes to spindle assembly during animal mitosis. J. Cell Biol. 167:831–840.

18. Kitamura, E., K. Tanaka, S. Komoto, Y. Kitamura, C. Antony, and T. U. Tanaka, 2010. Kinetochores Generate Microtubules with Distal Plus Ends: Their Roles and Limited Lifetime in Mitosis. Dev. Cell. 18:248–259.

19. Karsenti, E., J. Newport, and M. Kirschner, 1984. Respective roles of centrosomes and chromatin in the conversion of microtubule arrays from interphase to metaphase. J. Cell Biol. 99:47s–54s.

20. Sikirzhytski, V., V. Magidson, J. B. Steinman, J. He, M. L. Berre, I. Tikhonenko, J. G. Ault, B. F. McEwen, J. K. Chen, H. Sui, M. Piel, T. M. Kapoor, and A. Khodjakov, 2014. Direct kinetochore–spindle pole connections are not required for chromosome segregation. J. Cell Biol. 206:231–243.

21. Miller, P. M., A. W. Folkmann, A. R. R. Maia, N. Efimova, A. Efimov, and I. Kaverina, 2009. Golgi-derived CLASP-dependent microtubules control Golgi organization and polarized trafficking in motile cells. Nat. Cell. Biol. 11:1069–80.

22. Semenova, I., D. Gupta, T. Usui, I. Hayakawa, A. Cowan, and V. Rodionov, 2017. Stimulation of microtubule-based transport by nucleation of microtubules on pigment granules. Mol. Biol. Cell. 28:1418–1425.

23. Prosser, S. L., and L. Pelletier, 2017. Mitotic spindle assembly in animal cells: a fine balancing act. Nat. Rev. Mol. Cell Biol. 18:187–201.

24. Valdez, V. A., L. Neahring, S. Petry, and S. Dumont, 2023. Mechanisms underlying spindle assembly and robustness. Nat. Rev. Mol. Cell Biol. 24:523–542.

25. Goshima, G., F. Nédélec, and R. D. Vale, 2005. Mechanisms for focusing mitotic spindle poles by minus end-directed motor proteins. J. Cell Biol. 171:229–240.

26. Elting, M. W., C. L. Hueschen, D. B. Udy, and S. Dumont, 2014. Force on spindle microtubule minus ends moves chromosomes. J. Cell Biol. 206:245–256.

27. Sikirzhytski, V., F. Renda, I. Tikhonenko, V. Magidson, B. F. McEwen, and A. Khodjakov, 2018. Microtubules assemble near most kinetochores during early prometaphase in human cells. J. Cell Biol. 217:2647–2659.

28. Wollman, R., E. Cytrynbaum, J. Jones, T. Meyer, J. Scholey, and A. Mogilner, 2005. Efficient chromosome capture requires a bias in the ‘search-and-capture’ process during mitotic spindle assembly. Curr. Biol. 15:828–832.

29. Athale, C. A., A. Dinarina, M. Mora-Coral, C. Pugieux, F. Nedelec, and E. Karsenti, 2008. Regulation of microtubule dynamics by reaction cascades around chromosomes. Science. 322:1243–1247.

30. Carazo-Salas, R. E., G. Guarguaglini, O. J. Gruss, A. Segref, E. Karsenti, and I. W. Mattaj, 1999. Generation of GTP-bound Ran by RCC1 is required for chromatin-induced mitotic spindle formation. Nature. 400:178–181.

31. Paul, R., R. Wollman, W. T. Silkworth, I. K. Nardi, D. Cimini, and A. Mogilner, 2009. Computer simulations predict that chromosome movements and rotations accelerate mitotic spindle assembly without compromising accuracy. Proc. Natl. Acad. Sci. USA. 106:15708–15713.

32. Petry, S., A. C. Groen, K. Ishihara, T. J. Mitchison, and R. D. Vale, 2013. Branching microtubule nucleation in Xenopus egg extracts mediated by augmin and TPX2. Cell. 152:768–777.

33. Sánchez-Huertas, C., and J. Lüders, 2015. The Augmin Connection in the Geometry of Microtubule Networks. Curr. Biol. 25:R294–R299.

34. Kamasakim, T., E. O’Toole, S. Kita, M. Osumi, J. Usukura, J. R. McIntosh, and G. Goshima, 2013. Augmin-dependent microtubule nucleation at microtubule walls in the spindle. J. Cell Biol. 202:25–33.

35. Decker, F., D. Oriola, B. Dalton, and J. Brugués, 2018. Autocatalytic microtubule nucleation determines the size and mass of Xenopus laevis egg extract spindles. Elife. 7:e31149.

36. Gouveia, B., S. U. Setru, M. R. King, A. Hamlin, H. A. Stone, J. W. Shaevitz, and S. Petry, 2023. Acentrosomal spindles assemble from branching microtubule nucleation near chromosomes in Xenopus laevis egg extract. Nat. Commun. 14:3696.

37. Tulu, U. S., C. Fagerstrom, N. P. Ferenz, and P. Wadsworth, 2006. Molecular Requirements for Kinetochore-Associated Microtubule Formation in Mammalian Cells. Curr. Biol. 16:536–541.

38. Hayward, D., J. Metz, C. Pellacani, and J. G. Wakefield, 2014. Synergy between Multiple Microtubule-Generating Pathways Confers Robustness to Centrosome-Driven Mitotic Spindle Formation. Dev. Cell. 28:81–93.

39. Magidson, V., R. Paul, N. Yang, J. G. Ault, C. B. O’Connell, I. Tikhonenko, B. F. McEwen, A. Mogilner, and A. Khodjakov, 2015. Adaptive changes in the kinetochore architecture facilitate proper spindle assembly. Nat. Cell Biol. 17:1134–1144.

40. Travis, S. M., B. P. Mahon, and S. Petry, 2022. How Microtubules Build the Spindle Branch by Branch. Annu. Rev. Cell Dev. Biol. 38:1–23.

41. Murata, T., S. Sonobe, T. I. Baskin, S. Hyodo, S. Hasezawa, T. Nagata, T. Horio, and M. Hasebe, 2005. Microtubule-dependent microtubule nucleation based on recruitment of γ-tubulin in higher plants. Nat. Cell Biol. 7:961–968.

42. Verma, V., and T. J. Maresca, 2019. Direct observation of branching MT nucleation in living animal cells. J. Cell Biol. 218:2829–2840.

43. Chan, J., A. Sambade, G. Calder, and C. Lloyd, 2009. Arabidopsis cortical microtubules are initiated along, as well as branching from, existing microtubules. Plant Cell. 21:2298–2306.

44. Janson, M. E., T. G. Setty, A. Paoletti, and P. Tran, 2005. Efficient formation of bipolar microtubule bundles requires microtubule-bound γ-tubulin complexes. J. Cell Biol. 169:297–308.

45. Heald, R., R. Tournebize, T. Blank, R. Sandaltzopoulos, P. Becker, A. Hyman, and E. Karsenti, 1996. Self-organization of microtubules into bipolar spindles around artificial chromosomes in Xenopus egg extracts. Nature. 382:420–425.

46. Tängemo, C., P. Ronchi, J. Colombelli, U. Haselmann, J. C. Simpson, C. Antony, E. H. K. Stelzer, R. Pepperkok, and E. G. Reynaud, 2011. A novel laser nanosurgery approach supports de novo Golgi biogenesis in mammalian cells. J. Cell Sci. 124:978–987.

47. Ishihara, K., K. S. Korolev, and T. J. Mitchison, 2016. Physical basis of large microtubule aster growth. Elife. 5:e19145.

48. Mogilner, A., and A. Manhart, 2018. Intracellular Fluid Mechanics: Coupling Cytoplasmic Flow with Active Cytoskeletal Gel. Annu. Rev. Fluid Mech. 50:347–370.

49. Efimov, A., A. Kharitonov, N. Efimova, J. Loncarek, P. M. Miller, N. Andreyeva, P. Gleeson, N. Galjart, A. R. Maia, I. X. McLeod, et al., 2007. Asymmetric CLASP-dependent nucleation of noncentrosomal microtubules at the trans-Golgi network. Dev. Cell. 12:917–930.

50. Rivero, S., J. Cardenas, M. Bornens, and R. M. Rios, 2009. Microtubule nucleation at the cis-side of the Golgi apparatus requires AKAP450 and GM130. EMBO J. 28:1016–28.

51. Magidson, V., C. B. O’Connell, J. Lončarek, R. Paul, A. Mogilner, and A. Khodjakov, 2011. The Spatial Arrangement of Chromosomes during Prometaphase Facilitates Spindle Assembly. Cell. 146:555–567.

52. Nayak, P., S. Chatterjee, and R. Paul, 2023. Microtubule search-and-capture model evaluates the effect of chromosomal volume conservation on spindle assembly during mitosis. Phys. Rev. E. 108:034401.

53. Kalinina, I., A. Nandi, P. Delivani, M. R. Chacón, A. H. Klemm, D. Ramunno-Johnson, A. Krull, B. Lindner, N. Pavin, and I. M. Tolić-Nørrelykke, 2013. Pivoting of microtubules around the spindle pole accelerates kinetochore capture. Nat. Cell Biol. 15:82–87.

54. Blackwell, R., O. Sweezy-Schindler, C. Edelmaier, Z. R. Gergely, P. J. Flynn, S. Montes, A. Crapo, A. Doostan, R. McIntosh, M. A. Glaser, and M. D. Betterton, 2017. Contributions of Microtubule Dynamic Instability and Rotational Diffusion to Kinetochore Capture. Biophys. J. 112:552–563.

55. Tassin, A. M., B. Maro, and M. Bornens, 1985. Fate of microtubule-organizing centers during myogenesis in vitro. J. Cell Biol. 100:35–46.

56. Feldman, J. L., and J. R. Priess, 2012. A Role for the Centrosome and PAR-3 in the Hand-Off of MTOC Function during Epithelial Polarization. Curr. Biol. 22:575–582.

57. Ault, J. G., and C. L. Rieder, 1992. Chromosome malorientation and reorientation during mitosis. Cell Motil. Cytoskeleton. 22:155–159.

58. Lampson, M. A., and E. L. Grishchuk, 2017. Mechanisms to avoid and correct erroneous kinetochore-microtubule attachments. Biology. 6:1.

59. Lakshmi, R. B., P. Nayak, L. Raz, A. Sarkar, A. Saroha, P. Kumari, V. M. Nair, D. P. Kombarakkaran, S. Sajana, S. MG, et al., 2024. CKAP5 stabilizes CENP-E at kinetochores by regulating microtubule-chromosome attachments. EMBO Rep. 25:1909–1935.

60. Rusan, N. M., C. J. Fagerstrom, A.-M. C. Yvon, and P. Wadsworth, 2001. Cell Cycle-Dependent Changes in Microtubule Dynamics in Living Cells Expressing Green Fluorescent Protein-α Tubulin. Mol. Biol. Cell. 12:971–980.

61. Holy, T., and S. Leibler, 1994. Dynamic instability of microtubules as an efficient way to search in space. Proc. Natl. Acad. Sci. USA. 91:5682–5685.

62. Sarkar, A., R. Paul, and H. Rieger, 2019. Search and Capture Efficiency of Dynamic Microtubules for Centrosome Relocation during IS Formation. Biophys. J. 116:2079–2091.

