## Supplemental Information for "Branching, crosslinking and decentralization of microtubules accelerates intracellular assembly"

#### Supplemental Figures and Legends

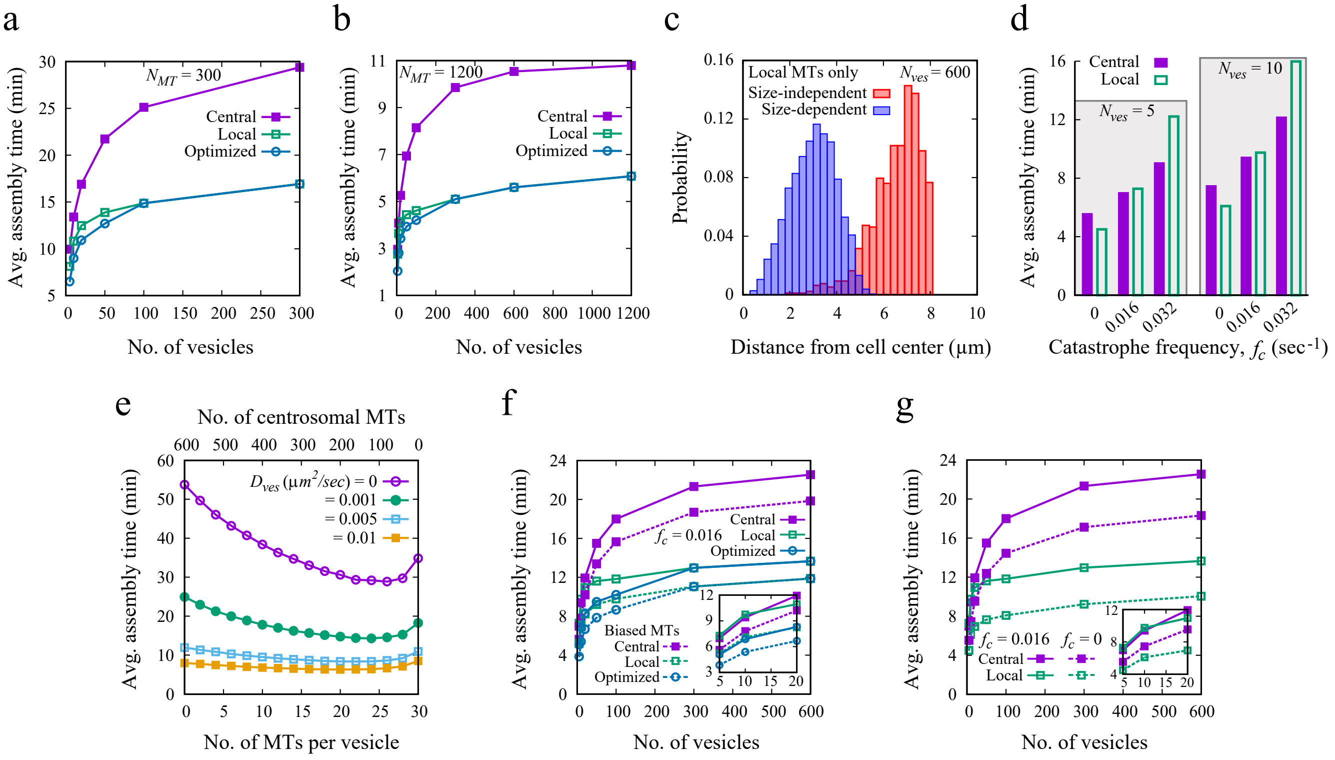


FIGURE S1 Effect of microtubule (MT) parameters and diffusion on vesicle assembly. (*a*–*b*) Assembly time for reduced ($N_{MT}$ = 300, *a*) and increased ($N_{MT}$ = 1200, *b*) total MTs, relative to the default ($N_{MT}$ = 600). MTs grow without spontaneous catastrophe ($f_c$ = 0) inside the cell. (*c*) The distance of the final vesicle aggregate from the cell center shows that, without central MTs, vesicles assemble within the cell interior, away from the center. Simulations were performed under two conditions: (i) Size-independent merging — the merged vesicle positions near either of the original vesicles, regardless of size (red bar), and (ii) Size-dependent merging — the merged vesicle settles near the larger one, resulting in a more inward position of the final aggregate (blue bar). Notably, average assembly times reported throughout the manuscript figures remain consistent across both scenarios. (*d*) Assembly time for $N_{ves}$ = 5 and 10 with MTs assigned to central or local searchers, shown for $f_c$ = 0, 0.016, and 0.032 sec$^{-1}$. (*e*) Assembly time decreases with increasing vesicle diffusion ($N_{ves}$ = 20). (*f*) Assembly time versus vesicle number for MT searchers assigned to central, local, or as per optimized combinations. Simulations use uniform catastrophe ($f_c$ = 0.016 sec$^{-1}$) and biased MTs ($d_{stab}$ = $R_c$/4, $\alpha$ = 10). (*g*) Comparison of uniform catastrophe ($f_c$ = 0.016 sec$^{-1}$) and no catastrophe. (*c*, *e*) Simulations use $f_c$ = 0.016 sec$^{-1}$.

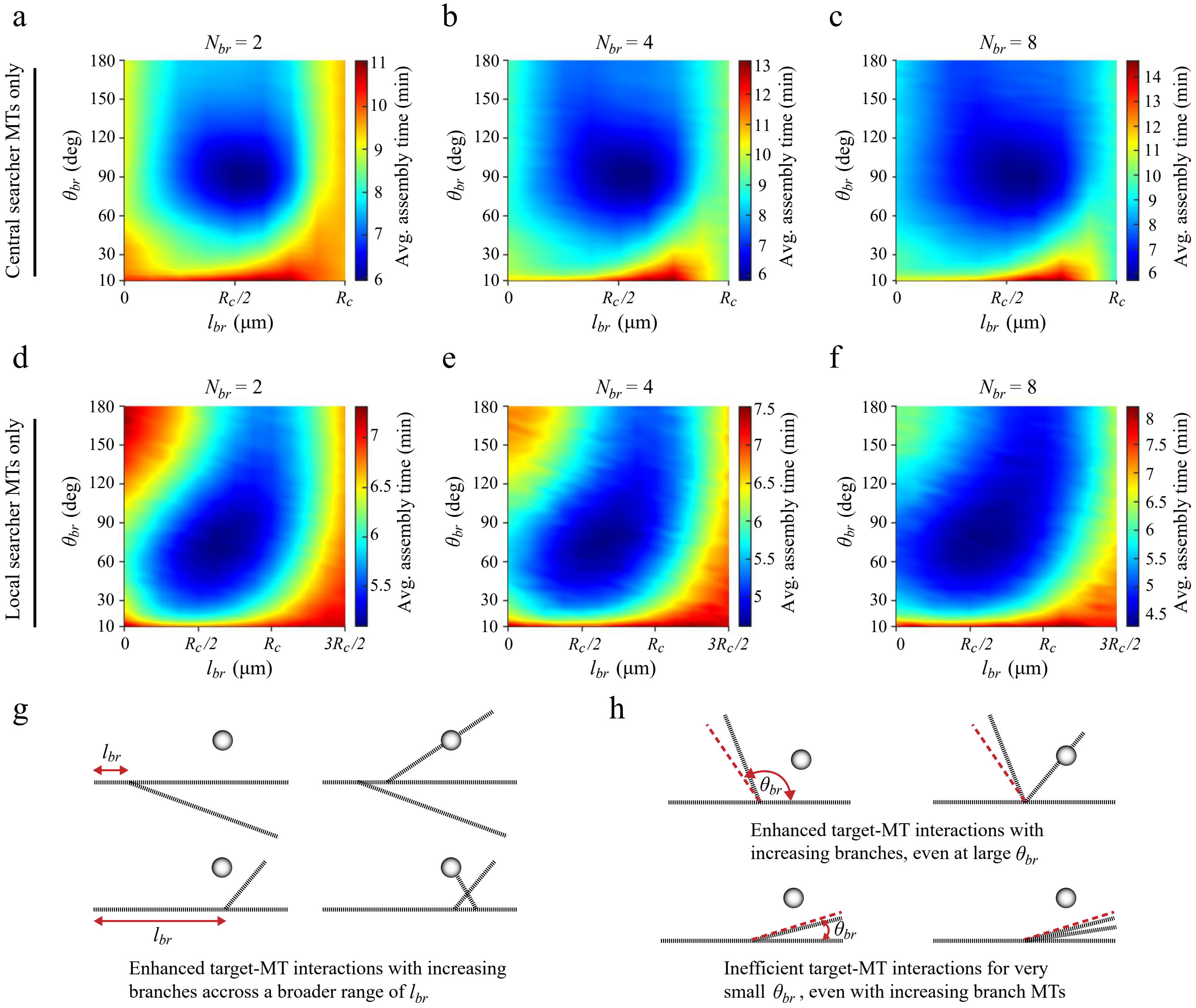


FIGURE S2 Vesicle assembly with varying numbers of branched MTs. (*a–c*) Assembly time versus branch length ($l_{br}$) and angle ($\theta_{br}$) for $N_{ves}$ = 20 with $N_{br}$ = 2, 4, and 8 MT branches, using only central searchers ($N_{cMT}$ = 600). (*d–f*) Similar analysis with all MTs as local searchers ($N_{vMT}$ = 600). (*g*) Schematic showing that increasing MT branches expands the search area, improving MT-vesicle encounters across a wider range of $l_{br}$. (*h*) At larger $\theta_{br}$, more branches enhance MT-vesicle interactions, while very small $\theta_{br}$ limits coverage and reduces assembly efficiency.

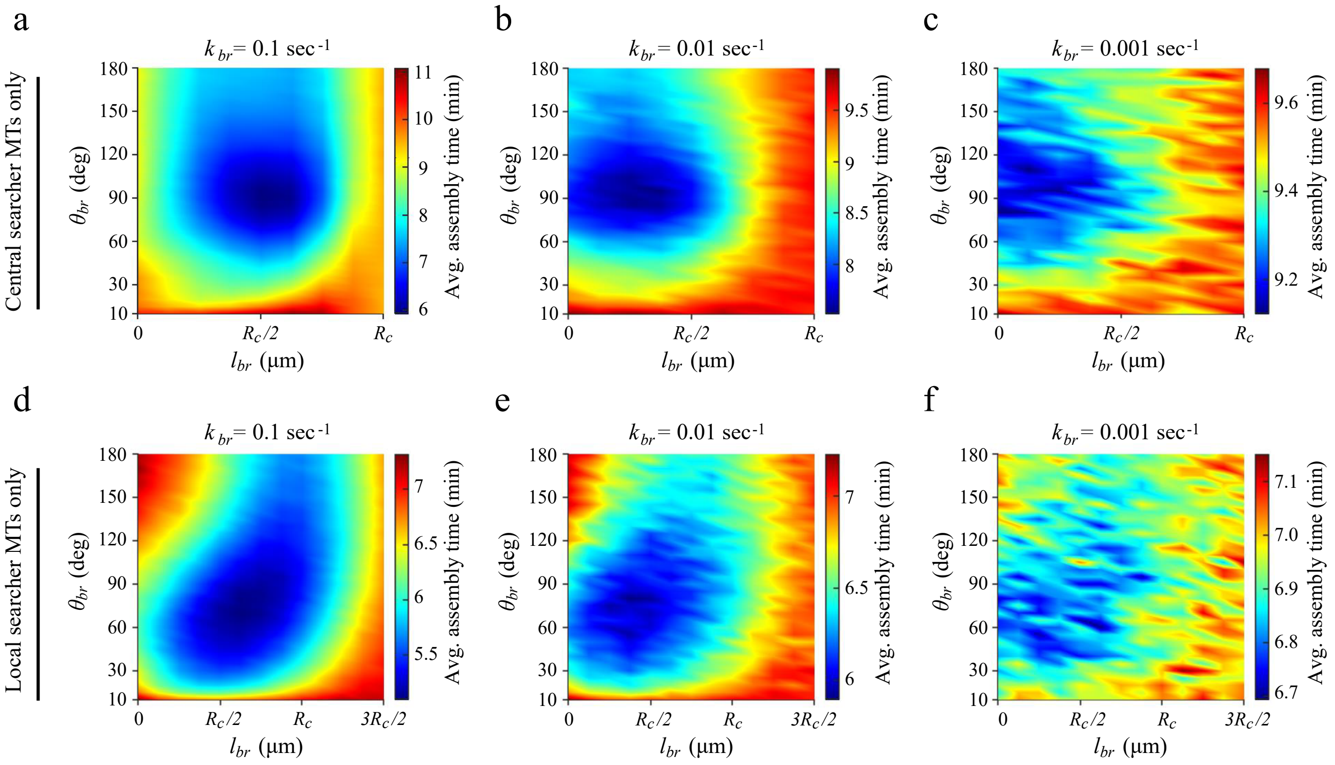


FIGURE S3 Vesicle assembly time across varying MT branching rates. Average assembly time as a function of $l_{br}$ and $\theta_{br}$ for $N_{ves}$ = 20 at branching rates $k_{br}$ = 0.1, 0.01, and 0.001 $sec^{-1}$, with MTs assigned to central searchers ($N_{cMT}$ = 600, *a*–*c*) or local searchers ($N_{vMT}$ = 600, *d*–*f*).

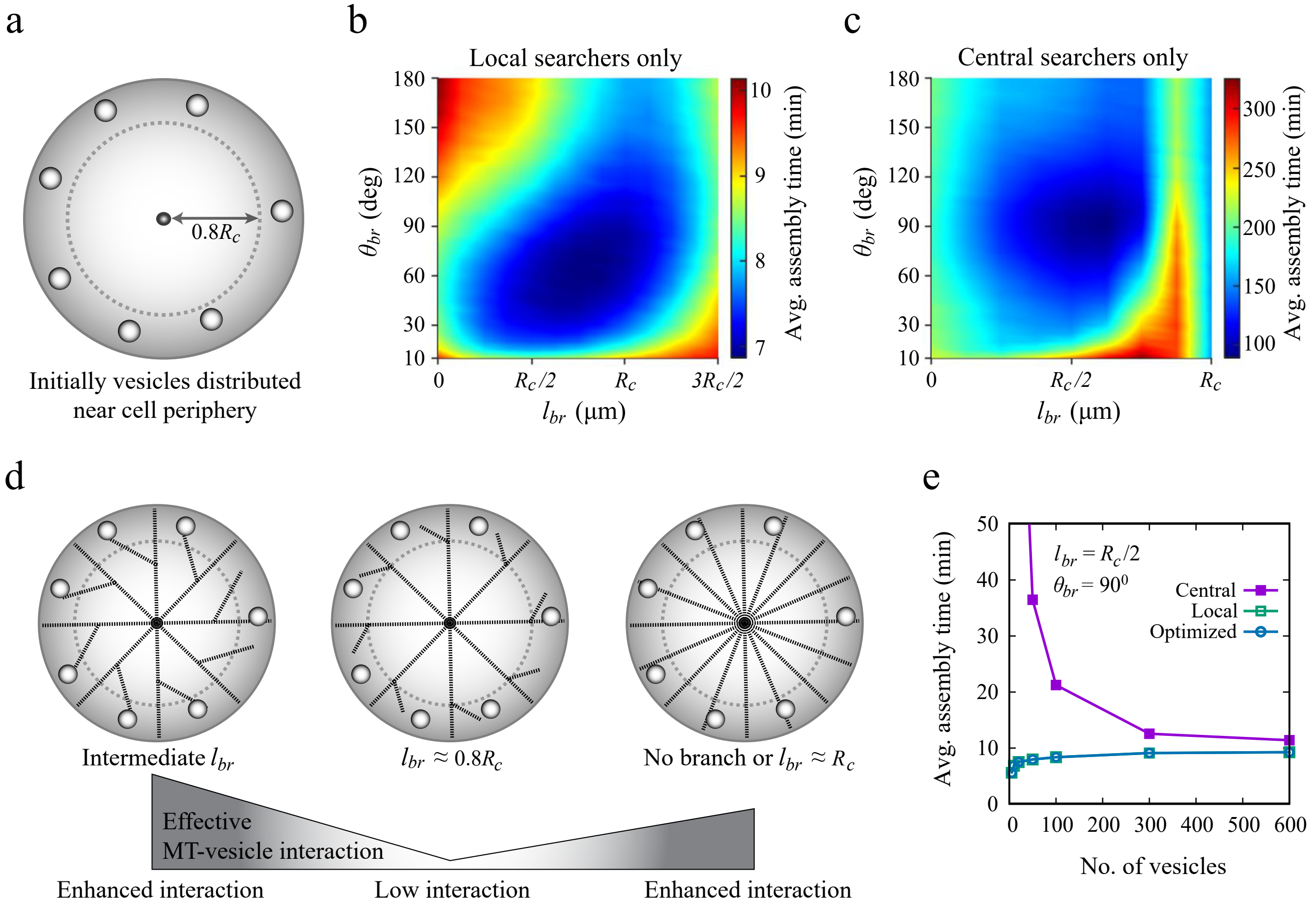


FIGURE S4 Vesicle assembly with peripheral vesicle distribution. (*a*) Vesicles are placed in an annular region of width $0.2R_c$ near the cell periphery. (*b*–*c*) Average assembly time as a function of branching length ($l_{br}$) and angle ($\theta_{br}$) for $N_{ves} = 20$, with MTs assigned to local searchers ($N_{vMT} = 600$, *b*) or central searchers ($N_{cMT} = 600$, *c*). (*d*) Schematic showing reduced MT-vesicle interactions for $l_{br} \approx 0.8R_c$ compared to intermediate ($l_{br} \approx R_c/2 \pm 2\mu$m) or larger values ($l_{br} \approx R_c$). For $l_{br} \approx R_c$, mother central searcher MTs grow close to the cell periphery, nullifying branching effects and aligning with the no-branching scenario. (*e*) Average assembly time vs. vesicle number for $l_{br} = R_c/2$ and $\theta_{br} = 90°$, comparing MTs exclusively assigned to central searchers, local searchers, or an optimized combination of both, yielding minimal assembly time.

### Supplemental Table

**TABLE S1 List of variables used in the model**

| Abbreviations | Meaning | Value \| Range \| Reference |
|---|---|---|
| $R_c$ | Cell radius | 10 µm |
| $R_{ves}$ | Initial vesicle radius | 0.25 µm |
| $N_{ves}$ | Number of initial vesicles | 5 – 600 |
| $N_{ch}$ | Number of chromosomes | 5 – 50 |
| $N_{kt}$ | Number of kinetochores | 10 – 100 |
| $R_{ch}$ | Chromosome radius | 0.7 µm |
| $l_{ch}$ | Chromosome length | 2 µm \| (1) |
| $R_{kt}$ | Kinetochore radius | 0.44 µm \| (1) |
| $l_{kt}$ | Kinetochore length | 0.35 µm \| (1) |
| $D_{ves}$ | Effective diffusion constant of the initial vesicles | 0.005 $\mu m^2/sec$ \| 0.001 – 0.01 $\mu m^2/sec$ |
| $D_{ch}$ | Effective diffusion constant of the chromosomes | 0.005 $\mu m^2/sec$ \| (1) |
| $N_{MT}$ | Total number of searcher MTs | 600 \| 300 – 1200 |
| $N_{br}$ | Number of MT branches per MT | 2 \| 2 – 8 |
| $k_{br}$ | Rate of branch initiation | 0.1 $sec^{-1}$\| 0.001 – 0.1 $sec^{-1}$ |
| $l_{br}$ | Cut-off distance for the branching MT nucleation | $0 - 3R_c/2$ |
| $\theta_{br}$ | Angle between the branch MTs and the mother central or local searcher MTs | $10° - 180°$ |
| $d_{stab}$ | Distance that determines the span of the stabilizing gradients centered around each vesicle | $R_c/8, R_c/4, R_c$ |
| $\alpha$ | A phenomenological constant determining the sensitivity of the catastrophe frequency to the stabilizing agents | 0 – 50 |
| $v_g$ | MTs growth velocity | 0.2383 $\mu m/sec$ \| (2) |
| $v_s$ | MTs shrinkage velocity | 0.2667 $\mu m/sec$ \| (2) |
| $f_c$ | MTs catastrophe frequency | 0 or $v_g.2/(3R_c)$ (0.016 $sec^{-1}$) \| (1, 3) |
| $f_r$ | MTs rescue frequency | 0 \| (3–5) |

### Supplemental Methods

#### Microtubule dynamics

Each microtubule (MT) is modeled as a cylindrical rod with a diameter of 25 nm and exhibits dynamic instability characterized by four parameters: growth velocity ($v_g$), shortening velocity ($v_s$), catastrophe frequency ($f_c$), and rescue frequency ($f_r$). During the growth phase, MTs elongate at velocity $v_g$ until a catastrophe occurs at rate $f_c$, initiating a shortening phase with velocity $v_s$. A nonzero rescue frequency $f_r$ can return a shortening MT to the growth phase. In our simulations, rescue events are excluded ($f_r = 0$), based on prior studies indicating that finite rescue increases search time by causing MTs to repeatedly grow in the same direction without a target there, prolonging futile searches (1, 3–5).

The dynamics of individual MTs were simulated using a Monte Carlo algorithm. At each time step $\Delta t$, a uniform random number between 0 and 1 was drawn and compared to the transition probability from growth to shortening, given by $1 - \exp(-f_c \Delta t)$. If the random number was below this threshold probability, the MT transitioned to the shortening state. Once an MT fully shortened back to the origin, a new growth cycle began, with the MT nucleating in a randomly chosen direction.

#### Introduction of bias in MT dynamics

To compute the spatial bias of the MT dynamics near the vesicles, we model the spatial gradient of a stabilizing factor as a linear superposition of exponential decay functions centered at each vesicle:

$$G_i = \sum_{k=1}^{N_{ves}} \frac{v_k}{v_{tot}} \exp\left(-\frac{d_{ik}}{d_{stab}}\right) \tag{S1}$$

Here, $v_{tot}$ is the volume of all vesicles, and $v_k$ is the volume of the $k$-th vesicle, so larger vesicles generate stronger gradients. $d_{ik}$ is the distance between the tip of the $i$-th MT and the center of the $k$-th vesicle. The decay length $d_{stab}$ determines how rapidly the influence of each vesicle decreases with distance.

The catastrophe frequency for the $i$-th MT then becomes:

$$f_c^i = f_c \exp(-\alpha G_i) \tag{S2}$$

Here, $f_c$ is the unbiased catastrophe frequency, and parameter $\alpha$ controls the MT's sensitivity to the stabilizing agents. As the value of parameter $\alpha$ increases, MTs become more sensitive to the bias, and the catastrophe frequency drops more steeply closer to the vesicles, vanishing right near the vesicles.

We apply the same framework to study the role of the RanGTP gradient in chromosomal assembly during mitosis.

#### Kinetics of Vesicles

Initially, small spherical vesicles are randomly distributed within the cellular volume. The position of each vesicle, $\mathbf{r}$ $(x_{ves}, y_{ves}, z_{ves})$, evolves over time $t$ according to

$$\frac{d\mathbf{r}(t)}{dt} = \sqrt{2D_{ves}}\,\boldsymbol{\eta}(t), \tag{S3}$$

where $D_{ves}$ is the diffusion coefficient of the vesicle. The components of the three-dimensional random vector $\boldsymbol{\eta}(t)$ $(\eta_1(t), \eta_2(t), \eta_3(t))$ follow a normal distribution with zero mean and unit variance. These components are statistically independent, satisfying $\langle \eta_i(t)\,\eta_j(t')\rangle = \delta_{ij}\,\delta(t-t')$, where $i, j = 1, 2, 3$, and $\delta_{ij}$ and $\delta(t-t')$ are the Kronecker and Dirac delta functions, respectively. Vesicle motion is confined to the interior of the cell by applying a reflecting boundary condition at the cell edge along the radial direction connecting the centers of the cell and vesicle.

#### Kinetics of Chromosomes

Initially, cylindrical chromosomes are randomly distributed within the cell, each with a random orientation. Orientation is defined using Euler angles $\phi_n$ $(n = 1, 2, 3)$, which allow full three-dimensional rotation of the chromosome along with its two associated cylindrical kinetochores. Chromosomes undergo translational diffusion characterized by a diffusion coefficient $D_{ch}$. We update their positions using an equation analogous to Eq. S3. Overlaps are avoided during position updates, and any movement beyond the cell boundary is restricted.

At each computational time step $\Delta t$, chromosomes also undergo rotational diffusion. This is modeled by applying small-angle rotations to the Euler angles:

$$\phi_n(t+\Delta t) = \phi_n(t) + d_{\text{rot}}\,\mathcal{N}(0,1), \quad n = 1, 2, 3, \tag{S4}$$

where $\mathcal{N}(0,1)$ denotes a normally distributed random variable with mean zero and unit variance, sampled independently for each $\phi_n$. The prefactor, $d_{\text{rot}} = 0.005$ radians, sets the typical magnitude of the rotational step.

#### Capture conditions

The successful capture of target vesicles (or kinetochores in mitotic spindle assembly) by searcher MTs from different origins can occur either through end-on attachment at the MT tip or lateral contact along the MT surface. According to the classical *search and capture* model, static targets can only be captured when the growing MT tip encounters them directly. In contrast, mobile targets such as vesicles or kinetochores typically form attachments through lateral interactions as they move within the cell and come into contact with existing MTs, often aided by motor proteins.

At each computational time step, end-on capture is evaluated by measuring the shortest distance between the MT tip and the center of the vesicle (or the axis of the kinetochore cylinder in chromosome models). If this distance is less than the radius of the target, capture is considered successful. For lateral capture, the shortest distance between the vesicle center (or kinetochore axis) and the MT axis is computed. A target is captured laterally if this distance is less than the sum of the MT and target radii.

In simulations that include MT–MT interactions, two MTs from different origins can capture each other at a rate $k_{\text{inter-MT}}$, provided their shortest inter-axial distance is less than the combined radii (each MT has a radius of approximately 12.5 nm). If an MT associated with a vesicle or kinetochore is captured in this way, the corresponding target vesicle or kinetochore is also considered captured.

#### Supporting References

1. Paul, R., R. Wollman, W. T. Silkworth, I. K. Nardi, D. Cimini, and A. Mogilner, 2009. Computer simulations predict that chromosome movements and rotations accelerate mitotic spindle assembly without compromising accuracy. *Proc. Natl. Acad. Sci. USA.* 106:15708–15713.

2. Rusan, N. M., C. J. Fagerstrom, A.-M. C. Yvon, and P. Wadsworth, 2001. Cell Cycle-Dependent Changes in Microtubule Dynamics in Living Cells Expressing Green Fluorescent Protein-$\alpha$ Tubulin. *Mol. Biol. Cell.* 12:971–980.

3. Wollman, R., E. Cytrynbaum, J. Jones, T. Meyer, J. Scholey, and A. Mogilner, 2005. Efficient chromosome capture requires a bias in the ‘search-and-capture’ process during mitotic spindle assembly. *Curr. Biol.* 15:828–832.

4. Holy, T., and S. Leibler, 1994. Dynamic instability of microtubules as an efficient way to search in space. *Proc. Natl. Acad. Sci. USA.* 91:5682–5685.

5. Sarkar, A., R. Paul, and H. Rieger, 2019. Search and Capture Efficiency of Dynamic Microtubules for Centrosome Relocation during IS Formation. *Biophys. J.* 116:2079–2091.
